# Single-cell and spatial atlas of steatotic liver disease-related hepatocellular carcinoma

**DOI:** 10.1101/2024.05.09.593073

**Authors:** Aldo Prawira, Hang Xu, Wei Qiang Leow, Masayuki Otsuka, Ziyao Chen, Mohammad Rahbari, Sharifah N. Hazirah, Martin Wasser, Alexander Chung, Brian K.P. Goh, Pierce K.H. Chow, Salvatore Albani, Joycelyn Lee, Tony K.H. Lim, Weiwei Zhai, Yock Young Dan, George Goh, David Tai, Ramanuj DasGupta, Mathias Heikenwalder, Jinmiao Chen, Valerie Chew

**Author notes:** **Correspondence:** Dr Valerie Chew, Translational Immunology Institute (TII), SingHealth-DukeNUS Academic Medical Centre, Singapore.

## Abstract

Steatotic liver disease-related hepatocellular carcinoma (SLD-HCC) poses significant challenges in liver cancer management. Our current study investigates the tumor microenvironment (TME) of SLD-HCC using single-cell transcriptomic, proteomic and spatial transcriptomic analyses. We identified altered immune-related and lipid metabolism pathways, particularly in regulatory T cells (Tregs) and cancer-associated fibroblasts (CAFs) within the SLD-HCC microenvironment, suggesting distinct cellular adaptations to a high-fat TME and general immunosuppression. Cytometry by time-of-flight revealed a cold and immunosuppressive TME depleted with CD8^+^ T cells and enriched with Tregs, while spatial transcriptomics uncovered a unique spatial architecture with Treg/CAF clusters specifically located at the tumor margins in SLD-HCC. Crucially, we identified Treg-CAF interactions as a key mediator associated with lack of response to immunotherapy in SLD-HCC. Our findings highlight the intricate immune dynamics of SLD-HCC, indicating potential therapeutic targets to counteract immune evasion and restore anti-tumor immunity in SLD-HCC.

**Highlights:**

- The TME of SLD-HCC exhibits altered immune and lipid pathways
- Immunosuppressive SLD-HCC TME is depleted with CD8^+^ T cells and enriched with Tregs
- SLD-HCC has a unique spatial architecture with Treg-CAF clusters at tumor margins
- Treg-CAF interaction via TNFSF14-TNFRSF14 axis mediates immunotherapy resistance

## Introduction

Hepatocellular carcinoma (HCC) is the third leading cause of cancer mortality worldwide, and its global incidence and mortality are predicted to rise by more than 50% by 2040^1^. HCC primarily develops from a background of chronic liver inflammation, with a heterogenous etiopathogenesis that includes chronic viral hepatitis infection, alcoholism and fatty liver diseases, making targeted therapy challenging^2^. Despite recent advancements in immunotherapies for advanced HCC, the objective response rate (ORR) remains modest, with 30% ORR for combination immunotherapy using atezolizumab (anti-PD-L1) and bevacizumab (anti-VEGFA)^3^. Since the majority of patients do not respond to the current immunotherapeutic strategies, a deeper understanding of the complex nature of the immune microenvironment in HCC is warranted.

Steatotic liver disease (SLD), encompassing alcoholic and non-alcoholic fatty liver disease (AFLD/NAFLD) or steatohepatitis (ASH/NASH), is detrimental to immune cell functions, resulting in immunosuppression and resistance to immune checkpoint inhibitors in HCC^4^. Recently, the Delphi consensus proposed the use of SLD as the overarching term to encompass the various aetiologies of hepatic steatosis, addressing the complex underlying disease conditions and social stigma associated with the terms “non-alcoholic” or “fatty”^5^. Furthermore, the challenge of retrospectively tracing alcohol intake in NAFLD patients who subsequently develop HCC underscores the need for a more inclusive terminology. The consensus also further redefined these SLD conditions to “metabolic dysfunction-associated steatotic liver disease” (MASLD) and “metabolic dysfunction-associated steatohepatitis” (MASH), respectively, to better reflect the underlying metabolic dysfunction. Given the rapid rise of SLD globally^6^, understanding the underlying mechanisms of SLD-related HCC (SLD-HCC) is crucial for the development of new immunotherapeutic strategies.

The early immune response to lipid accumulation in the liver was primarily mediated by myeloid subsets, particularly the Kupffer cells, which are the first line of defence against liver injury^7^. Additionally, XCR1^+^ dendritic cells drive inflammatory T cells reprogramming and progression to NASH^8^. The accumulation of PD-1^+^CD8^+^ T cells in the NASH liver, specifically the CXCR6^+^PD-1^+^ auto-aggressive and pro-inflammatory CD8^+^ T cells, has been associated with disease progression to HCC and poor response to checkpoint inhibition targeting PD-1/PD-L1 pathways in NASH-HCC^4,9^. Moreover, an increase in the regulatory T cell (Treg) population has been implicated in driving NASH-HCC through interaction with neutrophils, fostering an immunosuppressive microenvironment^10^. A recent study revealed an enrichment of exhausted PD-1^+^CD8^+^ T cells and PD-L1^+^ myeloid-derived suppressor cells in the TME of NASH-HCC^11^, providing insights into the immune microenvironment of NASH-HCC. However, comprehensive high dimensional single-cell and spatial analyses of tumor microenvironment (TME) in this HCC subtype remains underexplored.

Our present study focuses on SLD-HCC as the primary disease condition, leveraging on cutting-edge single-cell, spatial transcriptomics and proteomics analyses to unravel the complexity of the TME and anti-tumour immunity in SLD-HCC. Here, we show the unique cell populations that thrive and adapt to high-fat microenvironment. Importantly, we uncovered an interaction between Tregs and cancer-associated fibroblasts (CAFs) mediated by specific ligand-receptors, particularly at the margins of SLD-HCC tumors. This cellular interaction drives immunosuppression and is associated with the lack of response to immunotherapy. Taken together, our findings reveal the spatial immune-fibrotic interactome in SLD-HCC, with important implications for the understanding and management of this disease.

## Results

### Single-cell transcriptomic analysis of the SLD-HCC microenvironment reveals differential immune and lipid metabolism pathways

Using a multi-omics approach (Figure 1A and Table S1), we interrogated the TME of SLD-HCC (n=22) and non-SLD-HCC (n=31) patients, defined as ≥5% and <5% intrahepatic fat by pathological assessment, respectively. Additionally, we utilized tissue microarray from an independent cohort of 41 SLD-HCC and 62 non-SLD-HCC cases for validation using multiplex immunoflourescence (mIF). In concordance with the recent Delphi consensus^5^, all SLD-HCC cases in our cohort were also metabolic-associated SLD (MASLD) patients, as they harboured at least one metabolic risk factor (Table S1). Alongside steatosis, they also displayed signs of ballooning and inflammation marked by immune infiltration (Figure S1A). Comparison of key clinical parameters showed no significant differences that may confound the TME landscape or clinical outcomes between the two HCC subgroups (Figure S1B).

**Figure 1:**
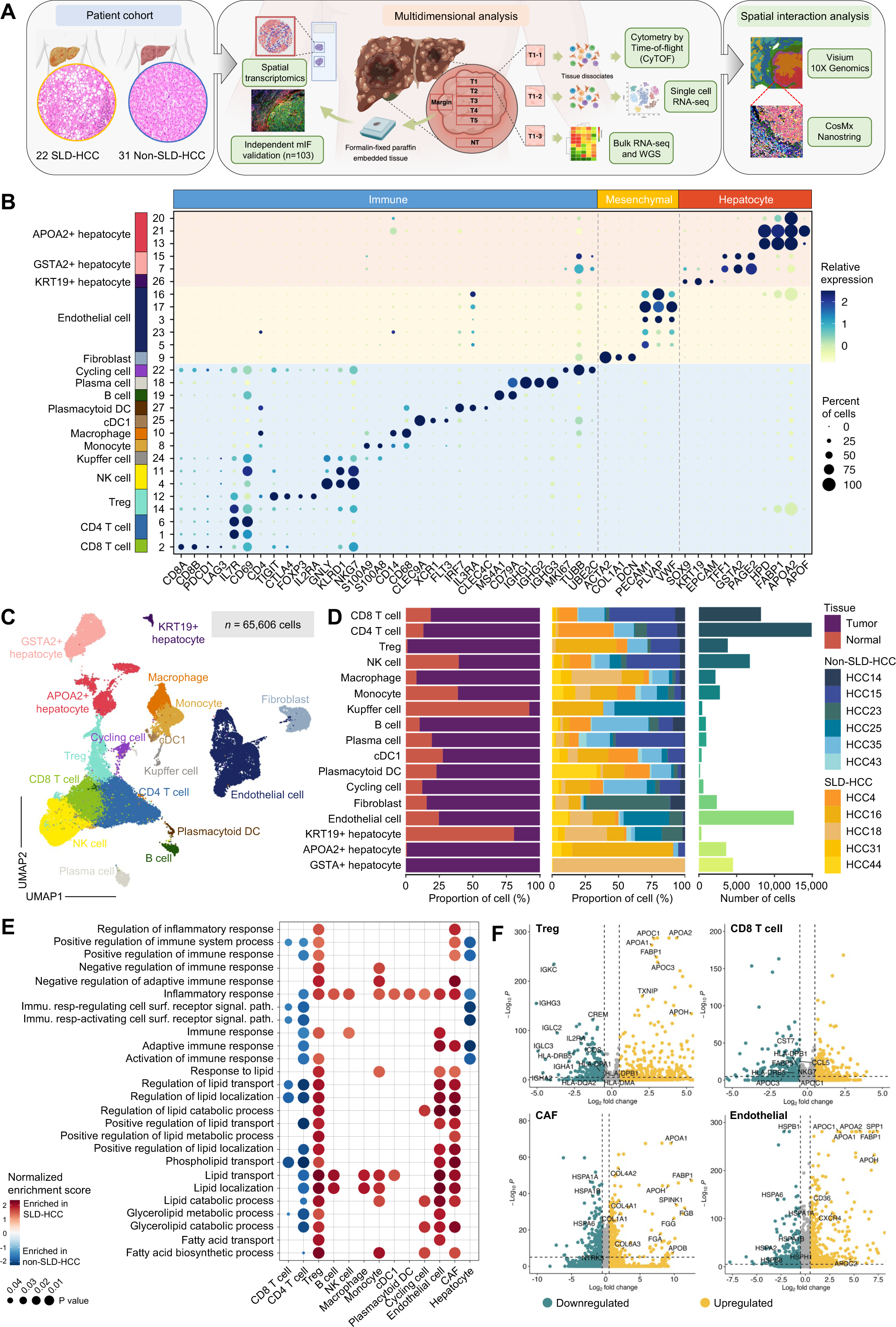
Single-cell transcriptomic landscapes of SLD- and non-SLD-HCC. **(A)** Schematic illustration of the study design. **(B)** Dot plot showing the top expressed genes in each of the 27 cell clusters identified from both tumors and non-tumor-adjacent liver tissues of patients with SLD-(n=5) and non-SLD-HCC (n=6). Dot color indicates average expression level, and dot size indicates the percentage of cells expressing the gene in each cluster. **(C)** UMAP plot showing Louvain clustering of 65,606 single cells from both tumors and non-tumor adjacent liver tissues of patients with SLD-(n=5) and non-SLD-HCC (n=6). **(D)** Bar graphs showing the distribution of major cell types between tumor/non-tumor tissues (left) and between patients with SLD-/non-SLD-HCC (middle) and the total numbers of each cell type (right). **(E)** Dot plot showing the enriched gene ontology pathways in each cell type, comparing differentially expressed genes in SLD- vs. non-SLD-HCCs. Dot colour indicates enrichment in SLD-compared to non-SLD-HCCs, and dot size indicates adjusted *p*-value as determined by Benjamini-Hochberg method. **(F)** Volcano plots depicting representative genes significantly up-(red) or downregulated (green) in Tregs, CD8^+^ T cells, fibroblasts and ECs from SLD-compared to non-SLD-HCCs. Differential gene expression analysis was conducted using the Wilcoxon rank sum test and log-transformed, with adjusted *p*-value < 0.05 considered significant. See also Figure S1 and Tables S1-S2.

First, we assessed the transcriptomic landscape of tumors and non-tumor sectors from five SLD-HCCs and six non-SLD-HCCs by performing single-cell RNA sequencing (scRNA-seq) analysis (Table S1 and STAR Methods). Using unsupervised clustering of 65,606 single cells, 27 clusters were identified based on the top expressed genes from each cluster (Figure S2A; Table S2 and STAR Methods). These clusters were further categorized into 17 major cell types according to similarities in their gene expression (Figures 1B–1C and S2B–S2C), encompassing immune, fibroblast, endothelial and hepatocyte populations originating from both tumor and non-tumor liver tissues. Cell proportion analysis showed that most hepatocytes were of tumor origin, while other cell types originated from both tumor and non-tumor tissues (Figure 1D). Tumor hepatocyte populations were heavily driven by individual patients, highlighting interpatient heterogeneity. In particular, *APOA2*^+^ hepatocytes mainly originated from SLD-HCCs (Figures 1D and S2C) and were enriched in lipid metabolism-related apolipoproteins^12^, consistent with the steatotic HCC phenotype. Other cell types were evenly contributed by different patients, reflecting consistencies in sample processing (Figures 1D and S2E–S2F).

Next, we isolated the cells from tumor tissues only and compared the phenotypes of individual cell types from SLD- vs. non-SLD-HCCs. Overall, pathway analyses suggested that both CD4^+^ and CD8^+^ T cells in the SLD-HCC TME downregulated pathways involved in immune or inflammatory responses (Figure 1E). Conversely, Tregs, endothelial cells (ECs) and CAFs showed upregulation of genes and cellular processes involved in lipid or fatty acid processes (Figure 1F), suggesting common adaptation to the steatotic microenvironment. Gene expression analysis of Tregs showed upregulation of genes involved in lipid metabolism including *TXNIP*, *FABP1* and members of the apolipoprotein family such as *APOA2*, *APOC1*, and *APOA1*^12^ (Figure 1F). This presumably provide Tregs with survival and functional advantages within the steatotic microenvironment. Conversely, genes associated with antigen presentation (*HLA-DRB5* and *HLA-DPA1*), lymphocyte activation (*CREM*, *IL2RA*, and *ICOS*), and immunoglobulin genes (*IGKC*, *IGHG3*, and *IGLC2*) were downregulated. In contrast, CD8^+^ T cells in SLD-HCC downregulated genes involved in both lipid metabolism (*FABP5*, *APOC3*, and *APOC1*) and cytotoxic immune function or antigen presentation such as *CST7*, *CCL5*, and *HLA-DPB1* (Figure 1F). Consistently, earlier studies have shown that lipid accumulation in the TME leads to CD8^+^ T cell exhaustion and dysfunction^13^.

Interestingly, CAFs in SLD-HCC upregulated pathways related to lipid metabolism (*APOA1*, *FABP1*, and *APOH*) (Figure 1F), suggesting a distinct metabolic reprogramming to meet the energy demands within the steatotic microenvironment. These fibroblasts also upregulated genes involved in fibrogenesis such as *COL4A2*, *COL4A1*, *FGB*, and *FGG* and downregulated heat-shock protein genes that are associated with regulation of fibrosis including *HSPA1A*, *HSPA1B*, and *HSPA6*^14^ (Figure 1F), suggesting their roles in fibrogenesis (Figure 1F). In SLD-HCC-associated ECs, we also observed similar upregulation of lipid metabolism genes (*FABP1*, *APOA1*, and *APOC1*) and *CD36*, known to regulate fatty acid uptake^15^ (Figure 1F). These ECs were also enriched in *CXCR4* and downregulated heat-shock protein genes (*HSPB1*, *HSPA6*, and *HSPA1A*) (Figure 1F), similar to the SLD-HCC-associated fibroblasts. Taken together, our scRNA-seq data suggest that a distinct metabolic reprogramming occurs within cells in the SLD-HCC microenvironment. Specifically, lipid metabolic pathways were enriched in Tregs, fibroblasts and ECs from SLD-HCCs, suggesting a shared adaptation to the steatotic microenvironment.

### Sub-clustering of T cells and stromal cells uncovers cellular diversity in SLD-HCC

To gain a deeper understanding of the cell populations enriched in lipid metabolism pathways in SLD-HCCs, we performed further characterization of T cells, fibroblasts and ECs sub-clusters. From a total of 26,192 T cells, unsupervised clustering yielded 14 sub-clusters (T1-T14) with distinct gene expression patterns originating from both tumor and non-tumor tissues, distributed between SLD- vs. non-SLD-HCCs (Figures 2A-2C and S3A–S3B; Table S3). We identified eight CD4-expressing (T1, T4, T6–T10, and T13) and four CD8-expressing (T2, T3, T5, and T11) T cell clusters (Figures 2D and S3C) as well as two mucosal-associated invariant T (MAIT) cell clusters (T12 and T14) that expressed signature genes including *SLC4A10*, *NCR3*, *TMIGD2*, *KLRB1* and *LST1*, previously reported in human solid tumors including HCC^16^ (Figures 2D and S3C). Among these clusters, the CD4^+^ T cell clusters T1, T6, and T8 were enriched in SLD-HCCs, while CD4^+^ T cell clusters T4 and T9, CD8^+^ T cell clusters T2, T3, and T11 and MAIT cluster T14 were enriched in non-SLD-HCCs (Figures 2B–2C).CD4-T1 and T6 resembled the Th1 phenotype, with higher expression of *NLRP3* and *LEF1* (Figure 2D). T6 was also enriched with heat-shock proteins (HSP) and inflammatory genes including *IFNG* and *TNF* (Figure 2D), indicating an enrichment of activated memory CD4^+^ T cells in SLD-HCCs. The other CD4^+^ clusters expressing *S1RP* (T4), *CCR4* (T10) or *ICOS* and *PDCD1* (T13) were either enriched in non-SLD-HCCs or evenly distributed between the HCC subtypes (Figures 2B-2C). There were also three *FOXP3*^+^*IKZF2*^+^ Treg clusters (T7, T8 and T9), with the SLD-HCC-enriched Treg-T8 showing higher expression of *FABP1* and apolipoproteins (Figure 2D) consistent with our previous observation (Figure 1G). Treg-T8 also resembled a lipid-associated Treg phenotype previously identified as a key contributor to the immunosuppressive TME and resistance to immunotherapy^17^. T7 and T9 both expressed canonical Treg genes such as *CTLA4*, *TIGIT* and *IL2RA* (Figure 2D). Meanwhile, most CD8^+^ T cell clusters were depleted from SLD-HCC TMEs (Figures 2B-2C). T3, T5 and T11 expressed *TOX*, with T3 and T11 expressing higher levels of *PDCD1* and *HAVCR2* (*TIM-3*) (Figure 2D), indicating exhausted phenotypes. Overall, we observed enrichment of activated memory CD4^+^ T cells (T1 and T6) and APO^+^ Tregs (T8) and a general depletion of CD8^+^ populations (T2, T3 and T11) in SLD-HCCs, indicating a more immunosuppressive and immunologically cold TME.

**Figure 2:**
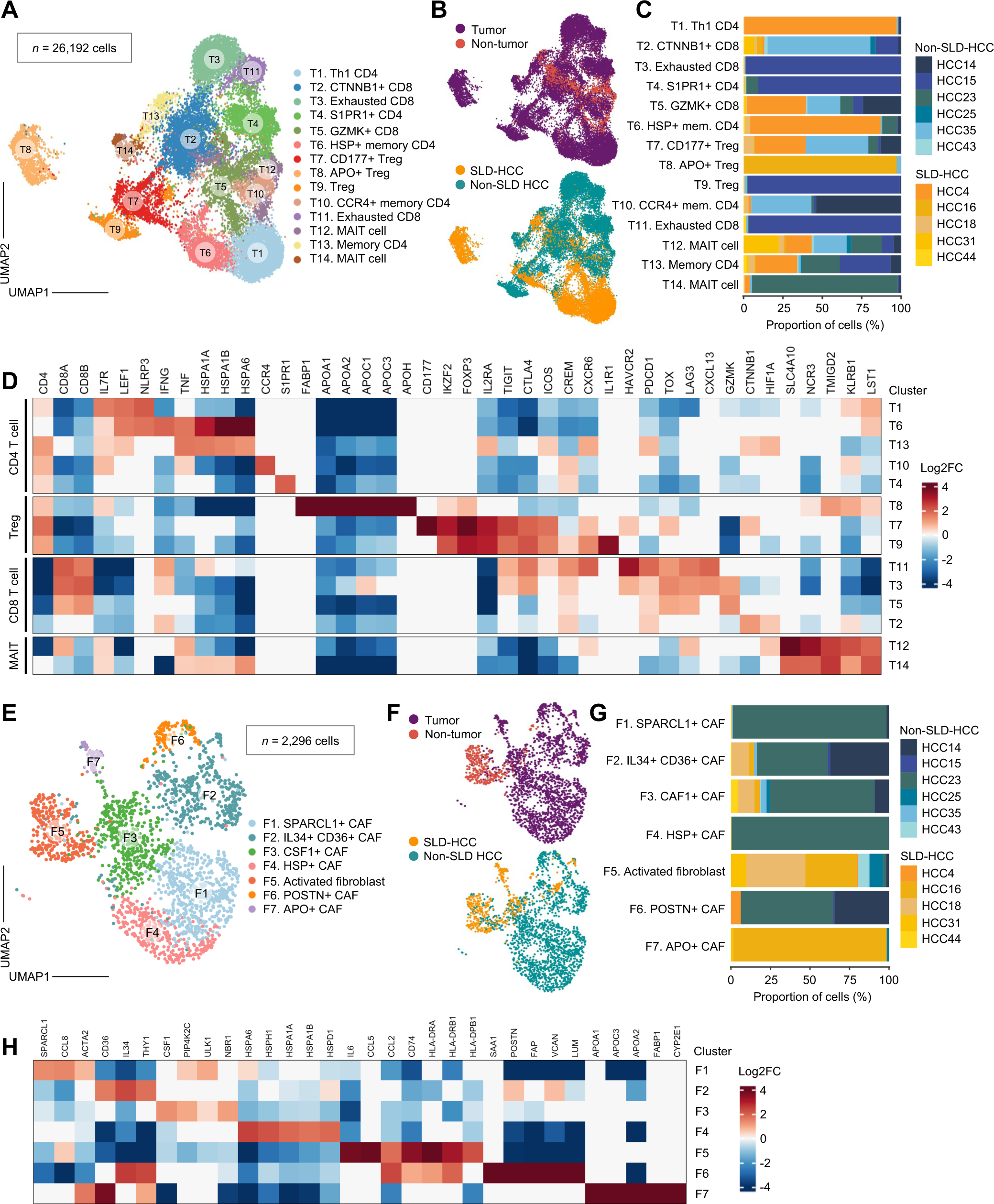
Characterization of T and mesenchymal cells in SLD-versus non-SLD-HCC. **(A)** UMAP plot showing Louvain clustering of 26,192 sub-clustered single T cells from tumors and non-tumor adjacent liver tissues of patients with SLD-(n=5) and non-SLD-HCC (n=6). **(B)** UMAP plot showing the distribution of T cells originating from tumor/non-tumor tissues (top) and from SLD- vs. non-SLD-HCCs (bottom). **(C)** Bar graphs showing the proportions of T cells within the 14 sub-clusters that originated from each SLD-/non-SLD-HCC patient. Each patient ID is colour-coded. **(D)** Heatmap showing the relative expression levels of cell type-specific genes in each T cell sub-cluster. **(E)** UMAP plot showing Louvain clustering of 2,296 sub-clustered single fibroblasts from tumors and non-tumor adjacent liver tissues of patients with SLD-(n=5) and non-SLD-HCC (n=6). **(F)** UMAP plot showing the distribution of fibroblasts originating from tumor/non-tumor tissues (top) and from SLD- vs. non-SLD-HCCs (bottom). **(G)** Bar graphs showing the proportions of fibroblasts within the seven sub-clusters that originated from each SLD-/non-SLD-HCC patient. Each patient ID is colour-coded. **(H)** Heatmap showing the relative expression levels of cell type-specific genes in each fibroblast sub-cluster. (D and H) The expression levels of genes in each cluster were compared against those of other clusters using the Wilcoxon rank sum test and log-transformed. The log_2_ fold change threshold were set to 0.25, and adjusted *p*-values of <0.05 were considered significant. See also Figures S2-S3 and Tables S3-S5.

Next, we further elucidated the fibroblast populations in SLD- and non-SLD-HCCs. We identified seven distinct sub-clusters (F1–F7) from 2,296 fibroblast cells (Figures 2E and S3D-S3E; Table S4), with both F5 and F7 enriched in SLD-HCCs (Figures 2F-2G). Interestingly, F5 displayed an activated or inflammatory phenotype with higher expression of *IL6*, *CCL5* and *CCL2* (Figure 2H). The F5 cluster also expressed *CD74* and antigen presentation genes such as *HLA-DRA*, *HLA-DRB1* and *HLA-DPB1* (Figure 2H), similar to a previously described antigen-presenting CAF that induces expansion of Tregs in pancreatic cancer^18^. Conversely, F7 expressed lipid metabolic genes including *FABP1*, *APOA1*, and *APOC3*^12^ (Figures 2F and 2G) as well as *CYP2E1*, involved in hepatic stellate cell activation and fibrosis^19^. Taken together, the CAF populations enriched in SLD-HCCs showed phenotypes implying involvement in remodelling of immune functions and activation of fibrosis. The remaining fibroblast clusters were almost exclusively present in non-SLD-HCC tumors (Figures 2F and 2G). For instance, *SPARCL1*^+^ CAFs (F1) represented a group of tumor vessel-associated fibroblasts related to reduced vascular invasion and superior survival in liver cancer patients^20^. *POSTN*^+^ CAFs (F6) showed higher expression of *FAP*, *POSTN*, and *LUM*, recently described as an important player in the oncofoetal system in HCC^21^.

Since ECs play a significant role in angiogenesis, tumor progression and metastasis and were enriched in lipid metabolism-associated genes in our dataset, we next sought to characterize these populations. A total of 10,347 ECs were clustered into 11 endothelial subtypes (E1–E11) based on their gene signatures (Figures S4A–S4D; Table S5). Of these, *PLVAP*^+^*HLA-DR*^+^ ECs (E4), pericytes (E10) and cycling ECs (E11) were enriched in SLD-HCCs, while *COL10A1*^+^ ECs (E2) and liver sinusoidal ECs (E8) were enriched in steatotic liver tissue (Figures S4E-S4F). In particular, *PLVAP*^+^*HLA-DR*^+^ ECs (E4) has been linked to immunomodulation within the onco-fetal ecosystem^22^. Interestingly, pericytes (E10) expressing characteristic genes such as *RGS5*, *PDGFRB* and *NOTCH3* were also enriched in SLD-HCCs (Figures S4C and S4F). Inflammation-driven fibrosis is known to induce the differentiation of pericytes to myofibroblast further exacerbating liver inflammation and injury^23^. We also found that liver sinusoidal ECs (LSECs; E8), characterised by the expression of *FCGR2B*, *CLEC1B*, *CLEC4G* and *LYVE1*, were enriched in steatotic liver tissue, (Figures S4C and S4F). These LSECs have been previously implicated in liver inflammation and progression of NASH^24^. In non-SLD-HCCs, we found enrichment of *PLPP3*^+^ (E1), *ACKR1*^+^ (E5), *IGFBP3*^+^ (E6) and *TMEM178A*^+^ ECs (E7) (Figures S4C-S4F). Overall, our findings demonstrated distinct T cell, fibroblast and EC populations within SLD- and non-SLD-HCCs, with SLD-HCC-enriched stromal cells displaying phenotypes related to immune remodelling and fibrosis.

### Single-cell proteomic immunoprofiling of SLD-HCCs reveals cold and immunosuppressive TME

As the results of our transcriptomic analyses indicated that major subsets enriched in SLD-HCCs were involved in immune modification, we further characterized the immune landscape of SLD-HCC using a single-cell proteomic approach. Here, tumor-infiltrating lymphocytes (TILs) from freshly resected tumor and non-tumor tissues were isolated and analysed via cytometry by time-of-flight (CyTOF; STAR Methods). In total, 1,663,336 single cells were analysed using a panel of antibodies targeting immune markers (STAR Methods). Unsupervised clustering using our in-house developed Extended Polydimensional Immunome Characterisation (EPIC) identified 49 clusters (Figures S5A–S5C), classified into major immune lineages including CD4^+^ T cells, CD8^+^ T cells, B cells and natural killer (NK) cells (Figure 3A).

**Figure 3:**
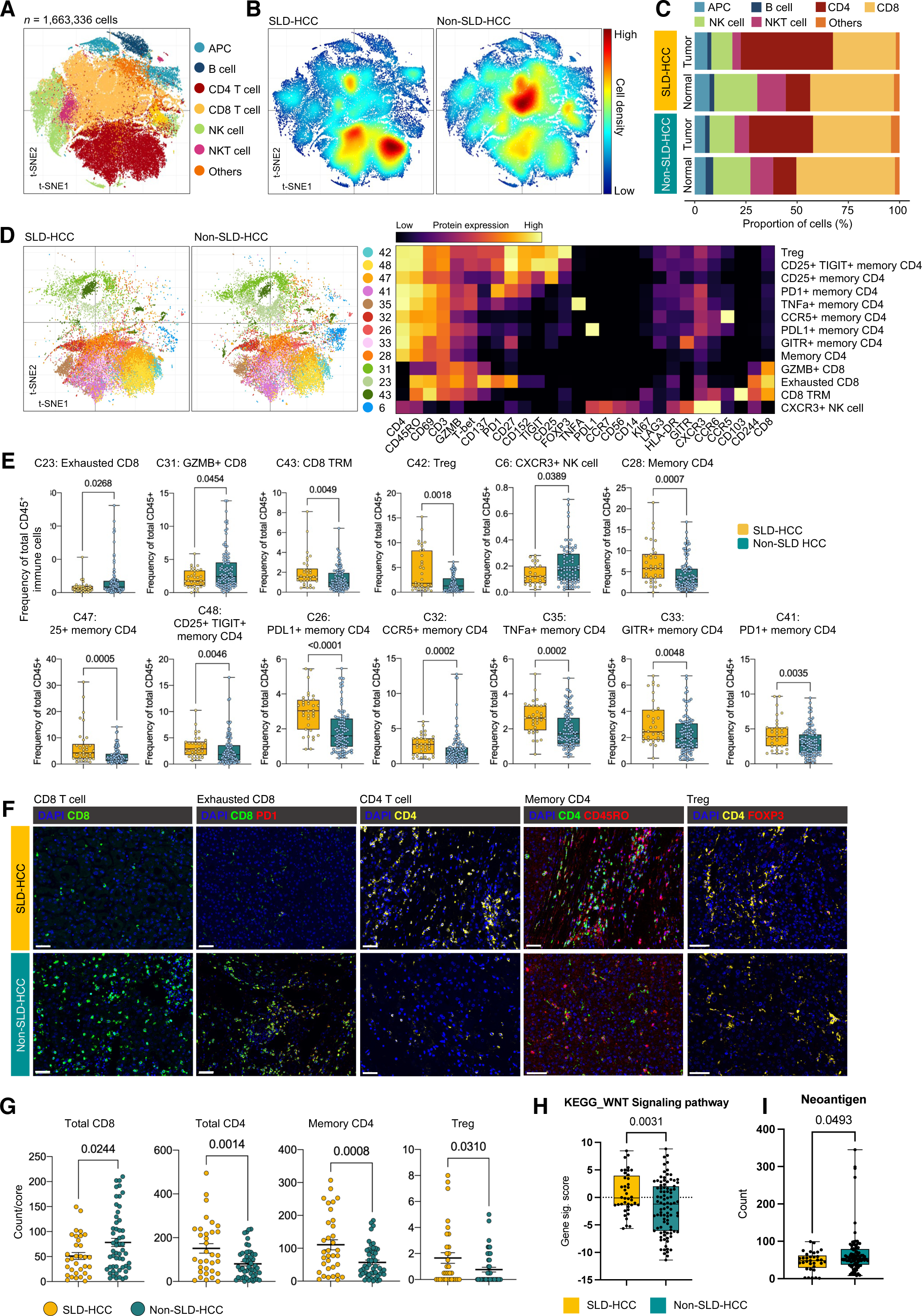
Immune landscapes profiling of SLD- and non-SLD-HCC. **(A)** t-SNE plot representing unsupervised clustering of 1,663,336 single immune cells from tumor and non-tumor tissues (SLD-HCC n=17; non-SLD-HCC n=27) into seven major immune lineages. **(B)** t-SNE plots showing the density of immune cells in SLD- and non-SLD-HCCs. **(C)** Bar graphs showing the proportions of each major immune lineage in normal and tumor tissues from SLD- and non-SLD-HCCs. **(D)** t-SNE plots showing 11 immune subtypes enriched in SLD- or non-SLD HCCs (left, 50,000 representative cells shown per plot). Heatmap showing the relative expression levels of immune marker proteins in each of these immune subtypes (right). **(E)** Bar graphs showing the frequencies of each immune subtype within total CD45^+^ cells from SLD- and non-SLD HCCs. *p*-values calculated by two-tailed Mann-Whitney test. **(F)** Representative multiplexed immunohistochemical staining of CD8^+^ T cells, exhausted CD8^+^ T cells, CD4^+^ T cells, memory CD4^+^ cells and Tregs in SLD- and non-SLD-HCCs from tissue microarrays comprising tumor cores from an independent cohort of 103 patients (SLD-HCC n=41; non-SLD-HCC n=62). Scale bar denotes 40 µm. **(G)** Graphs showing mean ± standard error of the mean (SEM) of total CD8, total CD4, memory CD4, and CD4^+^FOXP3^+^ Tregs in SLD- vs. non-SLD-HCCs. Cell density reported as count/core, each core= 0.785mm^2^. SLD-HCC n=41; non-SLD-HCC n=62. **(H)** KEGG_Wnt gene signalling scoring based on bulk RNA-seq data from tumor sectors from SLD-(n=40) and non-SLD-HCCs (n=91). **(I)** Neoantigen scores based on whole genome sequencing (WGS) from n=33 vs. n=90 tumor sectors from SLD- and non-SLD-HCCs, respectively. (**E, H and** I) Boxplots show median and the whiskers represent minimum and maximum values with the box edges showing the first and third quartiles. (**G-I**) *p*-values were calculated by two- tailed Student’s *t*-test. See also Figures S4-S5, Table S1 and STAR Methods.

In general, we observed a higher density of intratumoral CD4^+^ T cells and depletion of CD8^+^ T cells in SLD-HCCs compared to non-SLD-HCCs (Figures 3B–3C and S5D). Several previously reported immunosuppressive CD4^+^ T cells associated with poorer prognosis, including CD25^+^FOXP3^+^ Treg (C42) and CD4^+^ memory T cells expressing markers such as PD-L1, GITR and PD-1 (C28, 47, 48, 26, 32, 35, 33 and 41)^25,26^, were significantly enriched in SLD-HCCs (Figures 3D–3E). We also observed a depletion of exhausted and granzyme B-expressing active CD8^+^ T cells (C23 and C31, respectively) and a natural killer (NK) cell cluster (C6) in SLD-HCCs (Figures 3D–3E). Collectively, these data indicate that the TME of SLD-HCCs is immunosuppressive and “cold” or non-T cell inflamed, corroborating our scRNA seq findings. To determine whether this immune landscape was established in the early stages of SLD, we examined the T cell proportions in scRNA-seq data obtained from patients with healthy liver, early stages of NAFLD or late stages of NASH (Figures S6A-S6B). Indeed, we observed a trend of increased Tregs in NAFLD and depletion of CD8^+^ T cells in NASH compared to healthy livers (Figures S6A-S6B), suggesting that the immunosuppressive microenvironment could be induced in the early phases of SLD.

Next, we performed mIF (STAR Methods) on tissue microarrays comprising tumor cores from an independent cohort of 103 patients with SLD-HCC (n=41) or non-SLD-HCC (n=62) based on the same criteria of ≥5% intrahepatic fat. We consistently observed significantly more CD4^+^ T cells, specifically the memory CD4^+^ T cells and Tregs, and a reduction in CD8^+^ T cells in SLD-HCCs (Figures 3F–3G). As a non-T cell-inflamed TME, marked by depletion of intratumoural cytotoxic CD8^+^ T cells, has been linked to the Wnt signalling pathway^27^, we examined our bulk RNA-seq data (Table S1) and scored each sample for the KEGG_Wnt signalling pathway. Indeed, SLD-HCCs showed higher scores for the Wnt signalling pathway, consistent with immune-excluded tumors (Figure 3H). SLD-HCCs also displayed a lower neoantigen score (Figure 3I), in agreement with a previous report where the number of predicted MHC class I-associated neoantigens correlated with cytolytic activity^28^.

Overall, we demonstrated that SLD-HCCs are characterised by an immune-excluded microenvironment with depletion of cytotoxic CD8^+^ T cells and enrichment of immunosuppressive memory CD4^+^ T cells and Tregs.

### Spatial transcriptomic analysis identifies unique spatial architecture of SLD-HCC

Next, to unravel the distinct spatial organisation of key populations in SLD-HCCs as revealed by the single-cell transcriptomic and CyTOF analyses, we performed spatial transcriptomics on tissue samples from seven SLD-HCCs and five non-SLD-HCCs using Visium spatial gene assay (Figure 4A; STAR Methods). In total, we analysed spatial transcriptomics data from 126,099 spots across 12 tissue sections with a median depth of 9,698 UMI/spot and 3,708 genes/spot (Figure S7A). We carried out data integration using Harmony^29^ to normalise data across all tissue sections as shown by homogenous distribution of data across all patient samples in the UMAP (Figure 4B and Figure S7B). Since Visium is a spot-based transcriptomics assay, each spot, with a diameter of 55 µm, captures transcripts from multiple cells. Hence, we performed scoring using the top 100 genes from each cell type identified in our scRNA-seq data (Figures 1C and 4A) to decipher the cellular composition within each spot. Collectively, we found that spots with higher scores for immune cells also exhibited higher scores for fibroblasts and ECs, suggesting that they were in close proximity within the tissue (Figures 4C and S7C). In contrast, hepatocyte-associated spots showed lower scores for immune cells, fibroblasts and ECs (Figures 4C and S7C).

**Figure 4:**
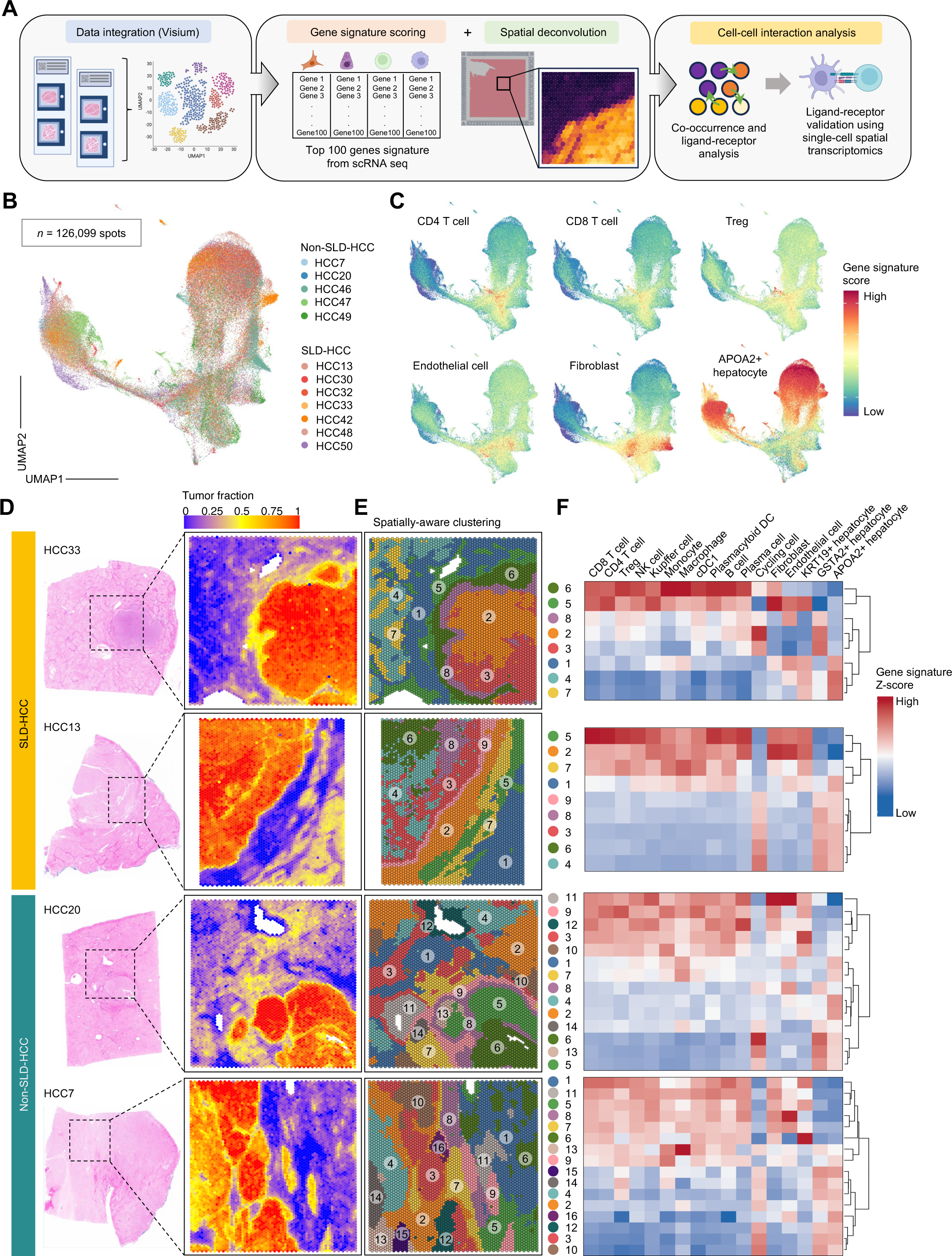
Spatial architecture of SLD- and non-SLD-HCC. **(A)** Data analysis workflow for Visium spatial transcriptomics (ST). Data integration was conducted using Harmony, and each spot was scored using the top 100 gene signatures from scRNA-seq using SpaCET. Cell-cell interactions were analysed using co-occurrence analysis. Ligand-receptor analysis was performed on spatially aware clustering data using Banksy, followed by L-R inference using NICHES. L-R interactions were further validated via COMMOT analysis. **(B)** UMAP plot illustrating the even distribution of ST data from patient samples after Harmony integration. n=126,099 spots from n=7 SLD-HCCs and n=5 non-SLD-HCCs. **(C)** UMAP plots showing gene signature scorings for major immune and non-immune subsets from all Visium data. **(D)** Representative H&E images on FFPE tissues comprising tumors, non-tumors and margin areas from patients with SLD- and non-SLD-HCC (left); spatial deconvolution using SpaCET, indicating tumor and non-tumor regions (right). Box, field of view (FOV)= 6.5 x 6.5 mm. **(E)** Spatially aware clustering using Banksy, showing the different domains within each tissue sample. **(F)** Heatmap showing relative gene signature scoring used to estimate the cellular composition within each domain. See also Figure S6, Table S1 and STAR Methods.

To understand the cellular interactions within their spatial context, we analysed individual tissue samples by combining SpaCET^30^ and Banksy^31^. SpaCET leverages the gene signatures from The Cancer Genome Atlas (TCGA) dataset to map the location of tumor and adjacent non-tumor hepatocytes, which is consistent with the histologic annotation based on haematoxylin and eosin (H&E) images (Figure 4D). Separately, we used the spatially-aware clustering algorithm Banksy to segment each tissue into different domains based on similarities in gene expression (Figure 4E). Using our previous scRNA-seq gene scoring methods, we annotated each Banksy domain and observed a higher score for immune cells in domains where the fibroblast score was higher, particularly at the domains lining the tumor margin of SLD-HCCs (Figure 4F). The SpaCET deconvolution method based on the TCGA dataset also consistently showed colocalization of fibroblast with immune cells at the tumor margin of SLD-HCCs (Figure S7E), therefore providing additional validation independent of our scRNA-seq data. Conversely, non-SLD-HCCs exhibited a distinct spatial organisation wherein domains with high scores for fibroblasts and immune cells were distributed across the tissue, rather than enriched around the tumor border (Figures 4D and S7D-S7E).

Taken together, the Visium spatial transcriptomics data showed potential interactions between immune and fibroblast populations, particularly at the margin or tumor edge in SLD-HCCs, based on the overlapped spots with simultaneous enrichment of their respective gene signatures.

### Cell-cell interaction analysis reveals Treg-fibroblast interaction at the SLD-HCC tumor margin

The spot-based Visium spatial transcriptomic analysis allowed us to identify potential interactions between cell populations, but its limited resolution necessitated a more refined assay to further specify the immune subset(s) interacting with fibroblasts. To address this, we first employed CellChat^32^ analysis on our scRNA-seq data, which demonstrated a relative increase in interaction strengths in Tregs and CD4^+^ T cells in SLD-HCC (Figure S8A). In contrast, the relative interaction strength was decreased in plasmacytoid DC, macrophages, Kupffer cells and CD8^+^ T cells within SLD-HCCs (Figure S8A). Although CellChat analysis lacked spatial context, the inferred interactions from scRNA-seq data led us to hypothesise that the immunosuppressive Treg and CD4^+^ T cell subsets could be interacting with the fibroblasts along the tumor edge in SLD-HCCs. To further validate these cellular interactions, we performed 1,000-plex fluorescence *in situ* hybridisation using a CosMx^TM^ Spatial Molecular Imager^33^, which allows spatial transcriptomic interrogation at a single-cell resolution, on two SLD- and two non-SLD-HCCs. Tissue sections were stained for CD45 (immune cells), CD3 (T cells), pan-cytokeratin (epithelial cells) and DAPI (nuclei) (Figure 5A). Across these four tissue sections, we selected 50 fields of view (FOVs) that encompassed tumor core, tumor margin and non-tumor regions (Figure S9A). Cell segmentation based on a machine learning algorithm (STAR Methods) identified 155,527 cells across all FOVs, and Leiden clustering classified these into 25 cell types, annotated based on their top expressed genes (Figures 5A-5C; Table S6). Overall, we detected 52,118,623 transcripts from 993 genes, with a mean of 270 transcripts per cell.

**Figure 5:**
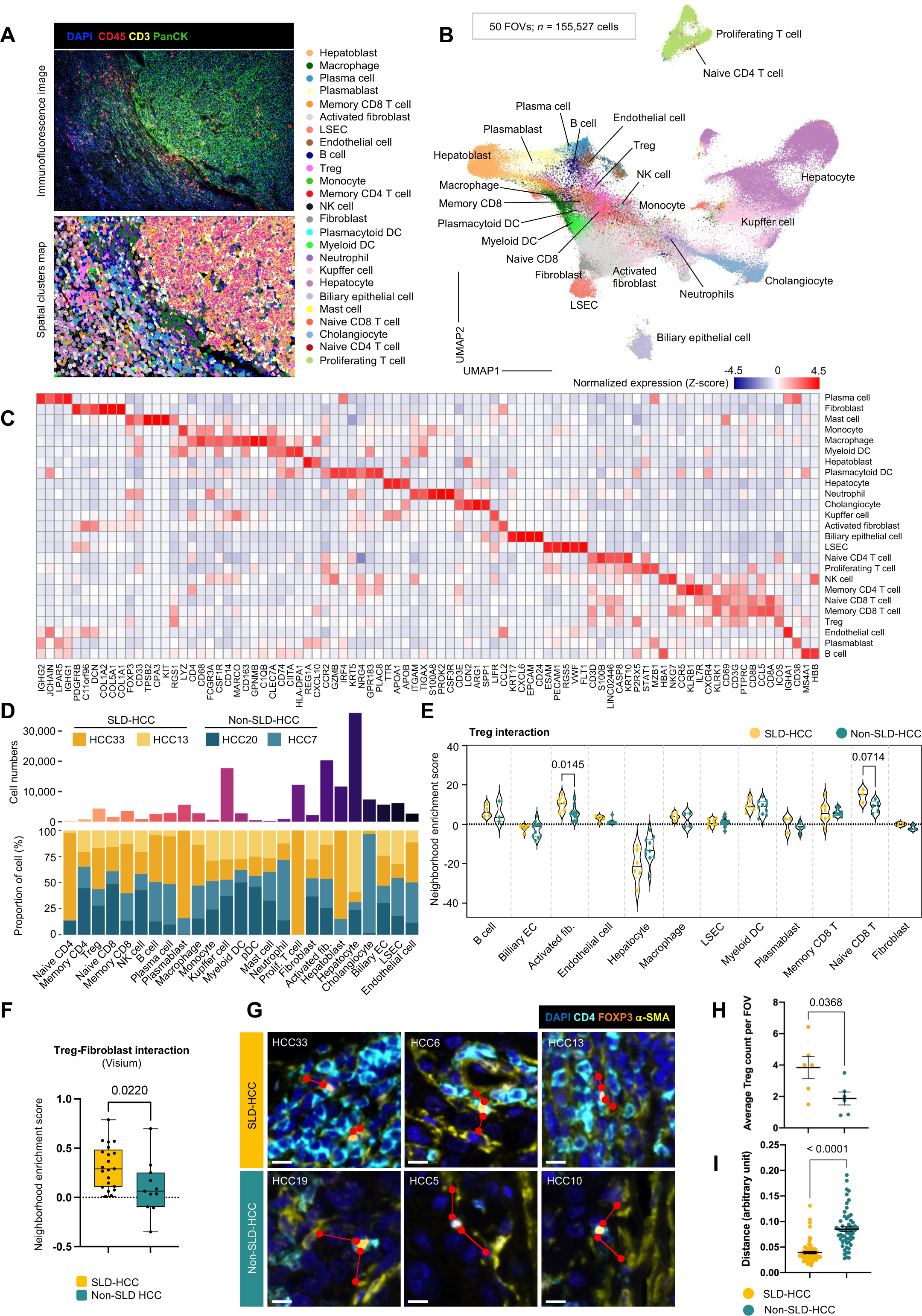
Cellular interaction network in SLD- and non-SLD-HCC. **(A)** Representative immunofluorescence (IF) image matched with ST data generated from CosMx unsupervised clustering. Each cluster is colour-coded accordingly. PanCK, pan-cytokeratin. Each FOV= 0.7 x 0.9mm. **(B)** UMAP plot illustrating all clusters from CosMx with n=155,527 total cells, n=50 FOVs from two SLD-HCCs and two non-SLD-HCCs. **(C)** Heatmap showing relative expression levels of selected genes representing each CosMx cluster. **(D)** Bar graphs showing the absolute number (top) and proportion within total cells (bottom) of each identified cell type across the two SLD-HCC and two non-SLD-HCC tissue samples. **(E)** Neighbourhood enrichment scores showing interaction strength between Tregs and other cell types from CosMx ST data. Note: only cell types shown in at least two FOVs per sample were included. Two-sided *p*-values calculated by pairwise Mann-Whitney test. **(F)** Neighbourhood enrichment scores showing interaction strength between Tregs and fibroblasts at margin domains from deconvoluted Visium ST data (n=7 SLD-HCCs and n=5 non-SLD-HCCs). Two-sided *p*-values calculated by pairwise Mann-Whitney test. **(G)** Representative IF images of margin areas from SLD- and non-SLD-HCC tissues stained for CD4, FoxP3 (Treg) and αSMA (fibroblast). DAPI was used for nuclear staining. Scale bar denotes 20 µm. **(H)** Comparison of mean number of Tregs between SLD- and non-SLD-HCC, quantified from 3- 5 randomly selected FOVs per tissue at tumor margin. **(I)** The distance between Treg and nearest fibroblast in the same FOVs from (H) were measured. The difference in distance between Treg and fibroblast were compared between SLD- and non-SLD-HCCs. (**F, H and I**) Boxplots show median and the whiskers represent minimum and maximum values with the box edges showing the first and third quartiles. (**H and I**) mIF data was obtained from six SLD- and six non-SLD-HCCs, and analysis was performed using Mann-Whitney U test. Graphs show mean ± standard error of the mean (SEM). See also Figure S7-S10, Table S6 and STAR Methods.

Immune and non-immune cell populations similar to our scRNA-seq clusters were identified, including CD4^+^ and CD8A^+^ T cells, myeloid and DC populations (*CD74*, *CIITA* and *HLA-DPA1*), plasma cells (*IGHG1*, *IGHG2* and *JCHAIN*) and fibroblasts (*COL1A1*, *COL1A2* and *COL5A1*) (Figures 5C and S10A; Table S6). Cell proportion analysis shows the distribution of cell populations across the four tissue samples, with hepatocytes being the most abundant cell type (Figure 5D). The Treg population expressed higher levels of *ICOS* and *PDCD1* (Figure 5C and Table S6), both of which are linked to a highly immunosuppressive Treg phenotype^34,35^. We also observed PD-1^+^ exhausted CD8^+^ T cells and CD163^hi^ tumor-associated immunosuppressive macrophages (Figure 5C and Table S6), validating the general immunosuppressive TME of HCC tumors. Of particular interest, we observed two distinct fibroblast populations; one was enriched with collagen genes, and the other resembled activated fibroblasts, expressing genes such as *CCL2*, *CXCL12* and *IL1R1* (Figure 5C and Table S6) that are important for immunosuppression and tumor progression^36^.

Next, we employed neighbourhood enrichment analysis on CosMx data to validate the potential cell-cell interactions identified in CellChat and Visium analyses. We first aimed to identify key immune cells potentially interacting with fibroblasts by focusing on the more abundant activated fibroblasts (Figure 5D) that resembled the SLD-HCC-enriched activated fibroblast cluster identified in scRNA-seq analysis (F5) (Figures 2F-2G). We found that this activated fibroblast cluster demonstrated significant interaction only with Tregs (Figure S11A). Reciprocally, we observed a significantly higher neighbourhood enrichment score between Tregs and activated fibroblasts and a trend toward increased interactions between Tregs and naïve CD8^+^ T cells in SLD-compared to non-SLD-HCCs (Figures 5E and S11B).

For cell-cell interaction analysis on the Visium data, we utilized Cell2location, which allowed higher sensitivity and resolution for fine-grained cell type deconvolution^37^ based on scRNA-seq data. Consistently, we found a significant correlation between Tregs and fibroblasts at the tumor margins in SLD- vs. non-SLD-HCCs (Figures 5F and S11C). To further validate this, we conducted mIF of Tregs and fibroblasts in six SLD-HCC and six non-SLD-HCC samples (Figure 5G). Because our previous data suggested that immune cells co-localise with CAFs along the tumor leading edge, we selected FOVs around the tumor border and quantified the abundance of Tregs and their distance from the nearest fibroblasts. Indeed, we found that the SLD-HCC tumor margins were enriched in Tregs (Figure 5H) and had a shorter average distance between Tregs and fibroblasts (Figure 5I), demonstrating the potential interaction between Treg and CAFs in the tumor leading edge in SLD-HCCs.

Taken together, our data indicate that Tregs, in addition to being more abundant (Figures 3E-3G) and having better adaptation (Figures 1E-1F) in the steatotic microenvironment, also show an increase in potential cell-cell interactions with fibroblasts at the margin of SLD-HCC tumors.

### Specific ligand-receptor pairs characterize Treg-fibroblast interactions in SLD-HCC

We next aimed to determine the ligand-receptor (L-R) pairs responsible for Treg-fibroblast communication by performing COMMOT (COMMunication analysis by Optimal Transport) analysis^38^ on the CosMx data. COMMOT is able to simultaneously analyse different L-R pairs, taking into consideration the spatial distances between specific cells from paired spatial data while also inferring spatial signalling directionality^38^. We identified a total of 11 common signalling pathways enriched in SLD-HCC, considering both Treg-fibroblast and fibroblast-Treg interactions (Figure 6A). To further validate these L-Rs, we also analysed the Visium data using NICHES (Niche Interactions and Communication Heterogeneity in Extracellular Signaling)^39^ to calculate the gene expression of L-R pairs for inferred interactions in each predefined domain from the spatially aware-clustering by Banksy (Figure 4D). Since we observed cell-cell interactions between fibroblasts and immune cells along the tumor margin, we isolated all clusters around the tumor margin (Figure 6B), which also exhibit higher expression of gene markers for fibroblasts and immune cells (Figure 4D). By comparing tumor margin clusters from SLD- vs. non-SLD-HCCs, NICHES analysis identified 12 L-R pairs that were exclusively expressed in the margins of SLD-HCC tumors (Figure S12A). Comparison between Treg-fibroblast-specific (Figure 6A) and margin-specific L-R targets (Figures 6B and S12A) revealed three common L-R pairs (Figure S12B). Among these, only TNFSF14-TNFRSF14 showed a significantly stronger COMMOT score in SLD-compared to non-SLD-HCCs (Figure S12C). Visualisation of TNFSF14-TNFRSF14 showed a spatial enrichment along the tumor leading edge in SLD-HCCs but further away from the tumor border in non-SLD-HCCs (Figures 6C and S12D).

**Figure 6:**
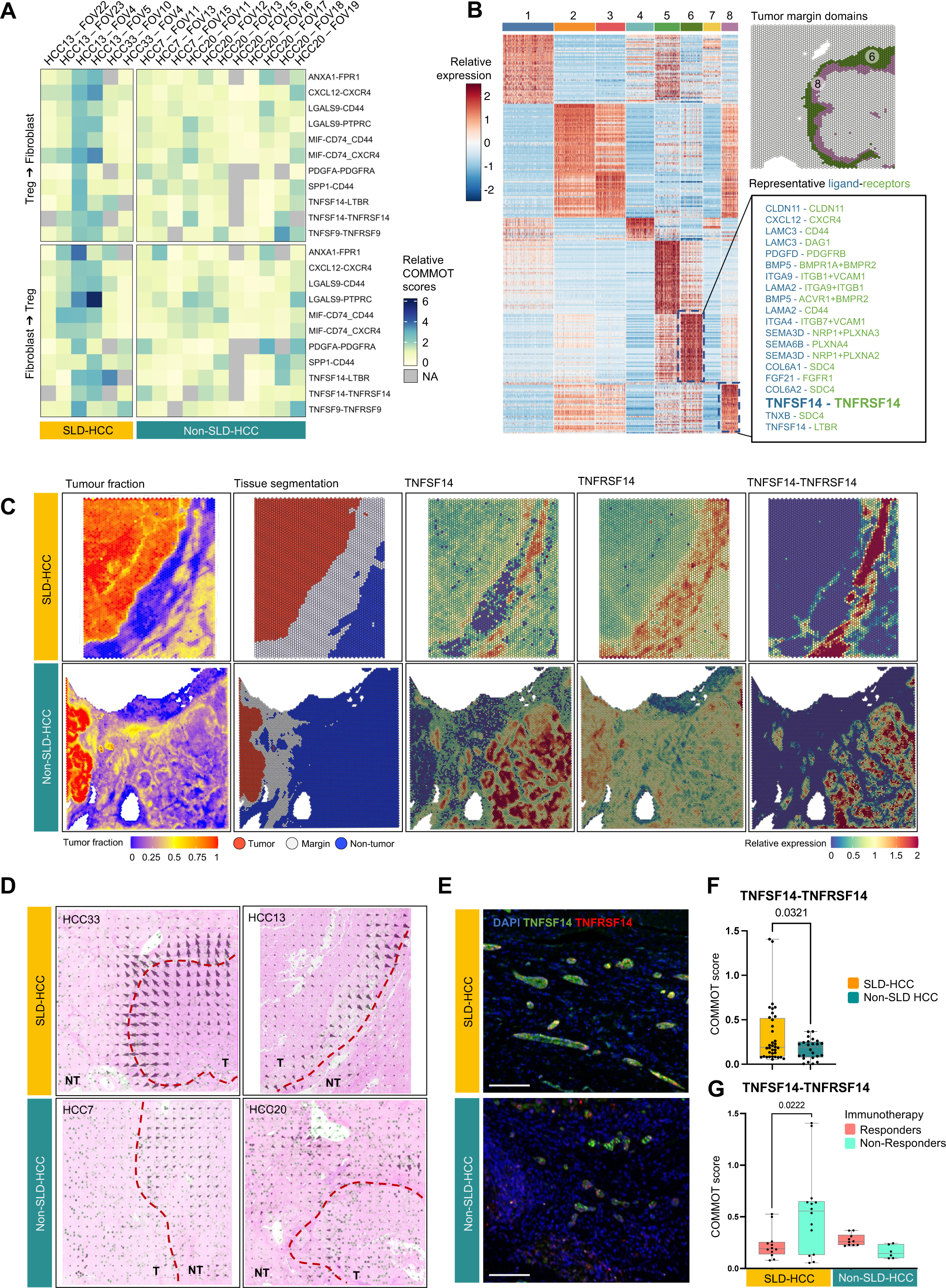
Ligand-receptor interactomes in SLD-HCC. **(A)** Heatmap showing relative COMMOT scores of enriched L-R pathways from CosMx data. **(B)** Heatmap showing relative expression levels of L-R pairs, determined by NICHES analysis on Visium ST data from SLD- vs. non-SLD-HCCs. Representative Visium map highlighting the tumour margin domains was shown (upper right). Specific enriched L-R pairs from clusters enriched at the tumor margin domains 6 and 8 were shown (Boxed, bottom right). **(C)** Representative images from Visium data showing tumor fraction scoring, tissue segmentation into tumor, margin and non-tumor regions as well as relative expression of TNFSF14-TNFRSF14 in SLD- vs. non-SLD-HCCs. **(D)** COMMOT analysis on Visium data showing distinct TNFSF14-TNFRSF14 strength and directionality in SLD- vs. non-SLD-HCCs. Gene expression intensity is marked by size and directionality by the pointed end of the arrows. Tumor (T) and non-tumor (NT) regions are separated by dashed red lines. **(E)** Representative IF images showing expression of TNFSF14-TNFRSF14 at tumor margins in SLD- and non-SLD-HCCs. Scale bar denotes 100 µm. **(F)** COMMOT scores comparing the strength of the TNFSF14-TNFRSF14 interaction between SLD-(n=7) vs. non-SLD-HCCs (n=5). Treg-Fib and Fib-Treg interactions from margin domains were analysed. **(G)** COMMOT scores comparing the strength of the TNFSF14-TNFRSF14 interaction between three responders and five non-responders to immunotherapy (pembrolizumab or atezolizumab+bevacizumab) in SLD-(n=5) vs. non-SLD-HCC (n=3). Treg-Fib and Fib-Treg interactions from margin domains were analysed. (**B-D**) Visium FOV= 6.5 x 6.5 mm. (**F and G**) Boxplots show median and the whiskers represent minimum and maximum values with the box edges showing the first and third quartiles. *p*-value determined by two-tailed Mann-Whitney test. See also Figure S11 and STAR Methods.

We further confirmed that TNFSF14-TNFRSF14 COMMOT signals showed a general directionality from tumor towards non-tumor areas (particularly at the tumor margin) in the SLD-HCC but random directionality in the non-SLD-HCC tumors (Figures 6D and S12E). mIF analysis of tumor margin areas validated this finding, showing enriched expression of TNFSF14-TNFRSF14 at the margin in SLD-HCCs (Figure 6E); these margins were previously shown to be enriched with Treg and fibroblast populations (Figures 5G-5I). Next, we performed COMMOT on the deconvoluted Visium spatial data based on our scRNA-seq database as described above. Consistently, we observed higher COMMOT scores for the TNFSF14-TNFRSF14 interaction specifically between Tregs and fibroblasts in SLD-compared to non-SLD-HCCs (Figure 6F). We also investigated the clinical significance of such interactions in HCC. We observed significantly higher COMMOT scores for TNFSF14-TNFRSF14 between Tregs and fibroblasts in the patients with SLD-HCCs who were non-responsive vs. those who were responsive to immunotherapy; this difference was not observed in the patients with non-SLD-HCCs (Figures 6G and S12F; Table S1). This finding suggests that the specific TNFSF14-TNFRSF14 interaction is associated with a lack of clinical response to immunotherapy in a subgroup of, but not all, patients with SLD-HCC.

## Discussion

Our current study comprehensively investigated the complex dynamics within the TME of SLD-HCC, employing a multi-omics approach to unravel its underlying molecular landscape and cell-cell interaction network. Through single-cell transcriptomic analyses, we uncovered an enrichment of lipid metabolic pathways particularly in Tregs, ECs and CAFs, indicating their adaptation to the steatotic liver microenvironment. Moreover, our single-cell proteomic immunoprofiling using CyTOF provided insights into the immune milieu of SLD-HCCs, indicating a predominantly cold (depleted with CD8^+^ T cells) and immunosuppressive (enriched with CD4^+^ memory T cells and Tregs) TME. This immunosuppressive phenotype, further characterized by dysregulated immune response pathways, underscores the formidable challenges in harnessing anti-tumor immunity within SLD-HCCs. Importantly, it’s noteworthy that a significant proportion of patients in our SLD-HCC cohort were also chronically infected with hepatitis viruses (Hep B or C). Despite this viral etiology, the observed immunosuppressive phenotype suggests dominant influence of the steatotic liver microenvironment. This distinction highlights a crucial aspect not typically addressed in previous studies, where patient stratification primarily focused on viral versus non-viral etiologies, often overlooking the potential impact of concurrent metabolic responses. Furthermore, leveraging recent advancements in spatial transcriptomics analysis that revolutionized our capability to better understand the TME within intact tumor tissues, we further unravelled the intricate spatial architecture of SLD-HCCs. This analysis revealed spatially segregated regions enriched with distinct cellular populations and signalling pathways. Notably, our investigation into Treg-fibroblast interactions uncovered key L-R pairs mediating crosstalk between these two critical populations, specifically at the tumor margin. The interaction between Tregs and fibroblasts was also linked to poor response to immunotherapy, shedding light on potential targets of therapeutic intervention specifically for SLD-HCCs.

The role of Tregs in maintaining immunosuppression within the TME and promoting tumor progression has previously been described in multiple cancers, including HCC^26^. However, their role in NASH remains controversial. One recent study reported an increase in Tregs in a NASH mouse model, but adoptive Treg transfer, which was intended for immunosuppression, unexpectedly worsened steatosis^40^. Another study described Tregs and neutrophils as key contributing players in driving carcinogenesis in NASH^10^, suggesting a complex role for Tregs in the progression of NASH to NASH-HCC. Indeed, our current data suggest that the SLD-HCC-enriched APO^+^ Tregs may function by interacting with fibroblasts in shaping the unique spatial architecture of SLD-HCC. This lipid-associated Treg population, previously described as an important population contributing to an immunosuppression in the TME and resistance to immunotherapy^17^, underscores the potential of targeting the lipid metabolic pathways within these cells for effective immunotherapy.

Conversely, fibroblasts play a central role in orchestrating liver fibrosis, a hallmark of advanced NASH. CAFs are thought to be actively involved in tumor progression through complex crosstalk with other cell types in the TME, and strategies targeting CAFs to augment immunotherapy have generated promising results in preclinical models and clinical trials^41^. A recent study in HCC also described a tumor immune barrier formed by interactions between CAFs and SPP1^+^ macrophages that limits immune infiltration and immunotherapy efficacy, highlighting the role of CAFs in immunomodulation of the TME^42^. However, little is known about the potential interactions between Tregs and CAFs in the context of NASH or NASH-driven HCC. One study in breast cancer reported that CAFs attract CD4^+^CD25^+^ T cells by producing CXCL12, retain them by OX40L, PPD-L1 and JAM2 and promote their differentiation to Tregs by inducing the expression of B7H3, CD73 and DPP4^43^. Here, we observed similar close interaction between Tregs and CAFs, with common molecular reprogramming marked by enhanced lipid metabolic pathways. This suggests a shared adaptation to the steatotic liver microenvironment, collectively driving the SLD-HCC disease phenotype.

TNFSF14 (Tumor Necrosis Factor Superfamily Member 14; also known as LIGHT), plays a crucial role in immunomodulation and inflammation in various diseases, including NASH^44^. Increased serum levels of TNFSF14 were observed in NAFLD patients^44^, indicating their potential role in the proinflammatory immune response. The roles of TNFSF14 and TNFRSF14 signalling in fibrosis^45^ and HCC^46^ have been previously described. Here, we show the interaction between Tregs and activated fibroblasts in SLD-HCCs via TNFSF14-TNFRSF14. The exact mechanism driving SLD towards HCC remains to be determined, and whether this interaction is critical or conserved during disease progression also warrants further study. More importantly, we demonstrated that Treg-CAF interactions via TNFSF14 and TNFRSF14 were associated with a lack of response to immunotherapy in HCC patients, suggesting that the response to immunotherapy is potentially dependent on the unique spatial architecture and interactome within the TME of SLD-HCCs.

Overall, our comprehensive multi-omics profiling of the SLD-HCC TME provides a holistic overview of the complex interplay between immune and stromal components in shaping tumor progression and therapeutic response. These findings deepen our understanding of SLD-HCC biology and offer valuable insights into potential avenues for targeted therapy and immunomodulation. We anticipate that our study will serve as a foundation for future investigations aimed at unravelling the intricacies of intratumoral cellular interactomes and facilitating the development of precision therapeutic strategies for patients with SLD-HCC.

## Acknowledgments

The authors would like to thank all members of TII, all participating patients, and the clinical research coordinators from NCCS, SGH and NUHS for their contributions to the patient sample collection.

This work was supported by the National Medical Research Council (NMRC), Singapore (reference number: CIRG22jul-0025, NMRC/TCR/015-NCC/2016, NMRC/CSA-SI/0013/2017, NMRC/CSA-SI/0018/2017, NMRC/OFLCG/003/2018), Duke-NUS Khoo Bridge Funding Award (Duke-NUS-KBrFA/2022/0058), International Gilead Sciences Research Scholars Program in Liver Disease – Asia and National Research Foundation, Singapore (ref number: NRF-CRP26-2021-0005).

## Author contributions

AP and HX: Investigation, formal analysis, visualization and writing. WQL, MO, ZC, MR, SNH and MW: Investigation and formal analysis. AC, BKPG, PKHC, SA, JL, TKHL, WZ and YYD: Resources and supervision. GG, DT, RDG, MH and JC: Resources, investigation and supervision. VC: Conceptualization, resources, formal analysis, supervision and writing.

