## Supplementary figures for "Single-cell and spatial atlas of steatotic liver disease-related hepatocellular carcinoma"

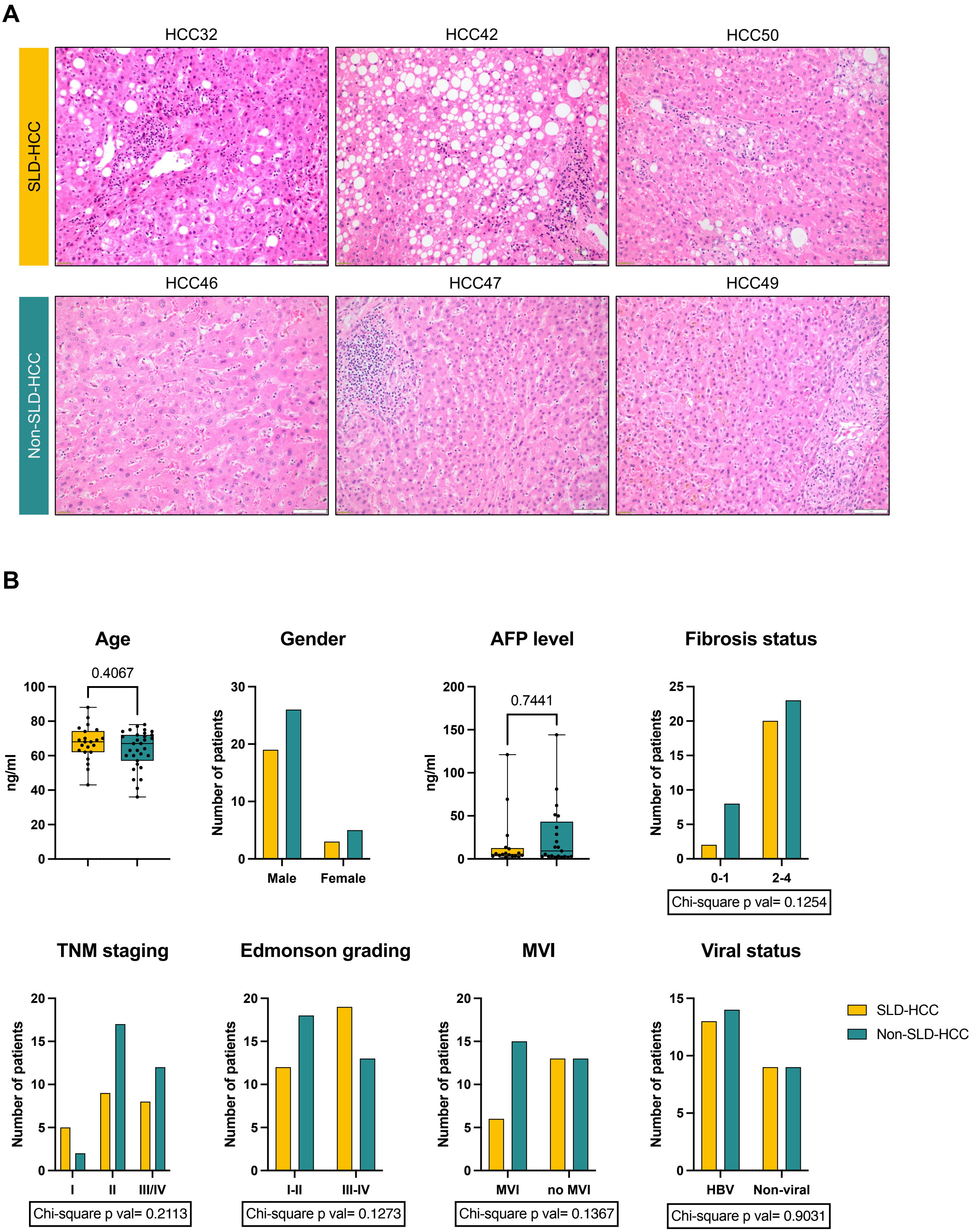


**Figure S1: Patient characterization for SLD- vs non-SLD-HCCs.**

**A.** Representative H&E images showing steatosis, ballooning and immune infiltration (inflammation) for SLD-HCC vs the lack of such features in non-SLD-HCC. Scale bar= 100 µm. **B.** Comparison of key clinical parameters between SLD-HCC vs non-SLD-HCCs.

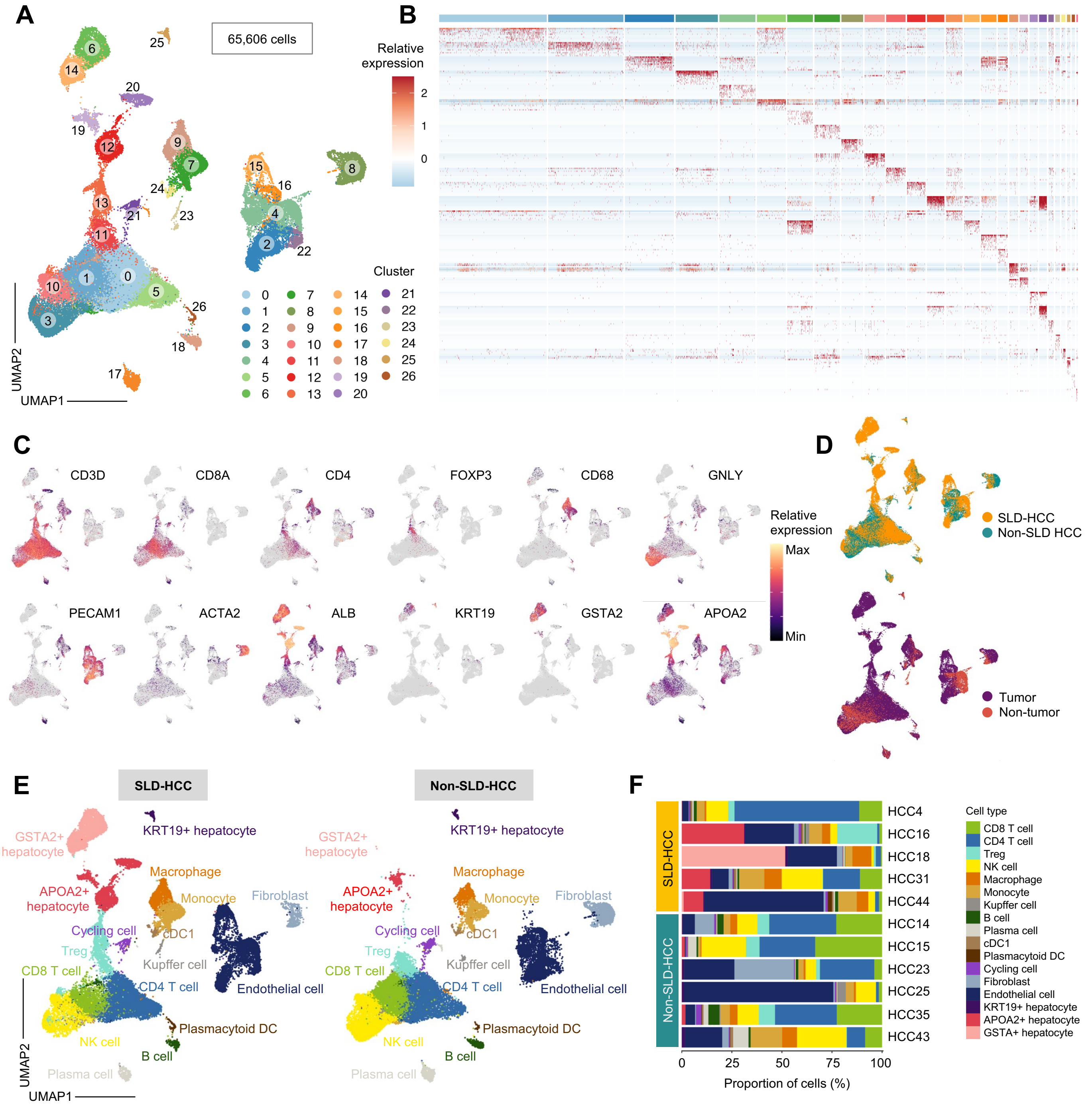


**Figure S2: Single cell RNA sequencing analysis of SLD- and non-SLD-HCCs.**

**A.** UMAP showing 27 clusters from 65,606 cells. **B.** Heatmap showing the top differentially expressed genes for each cluster. **C.** UMAP of selected genes for each cell type. **D.** UMAP showing distribution of cells according to disease status (upper) and tissue of origin (lower). **E.** UMAP showing individual clusters in SLD- and non-SLD-HCC, respectively. **F.** Cell proportion for each individual patient.

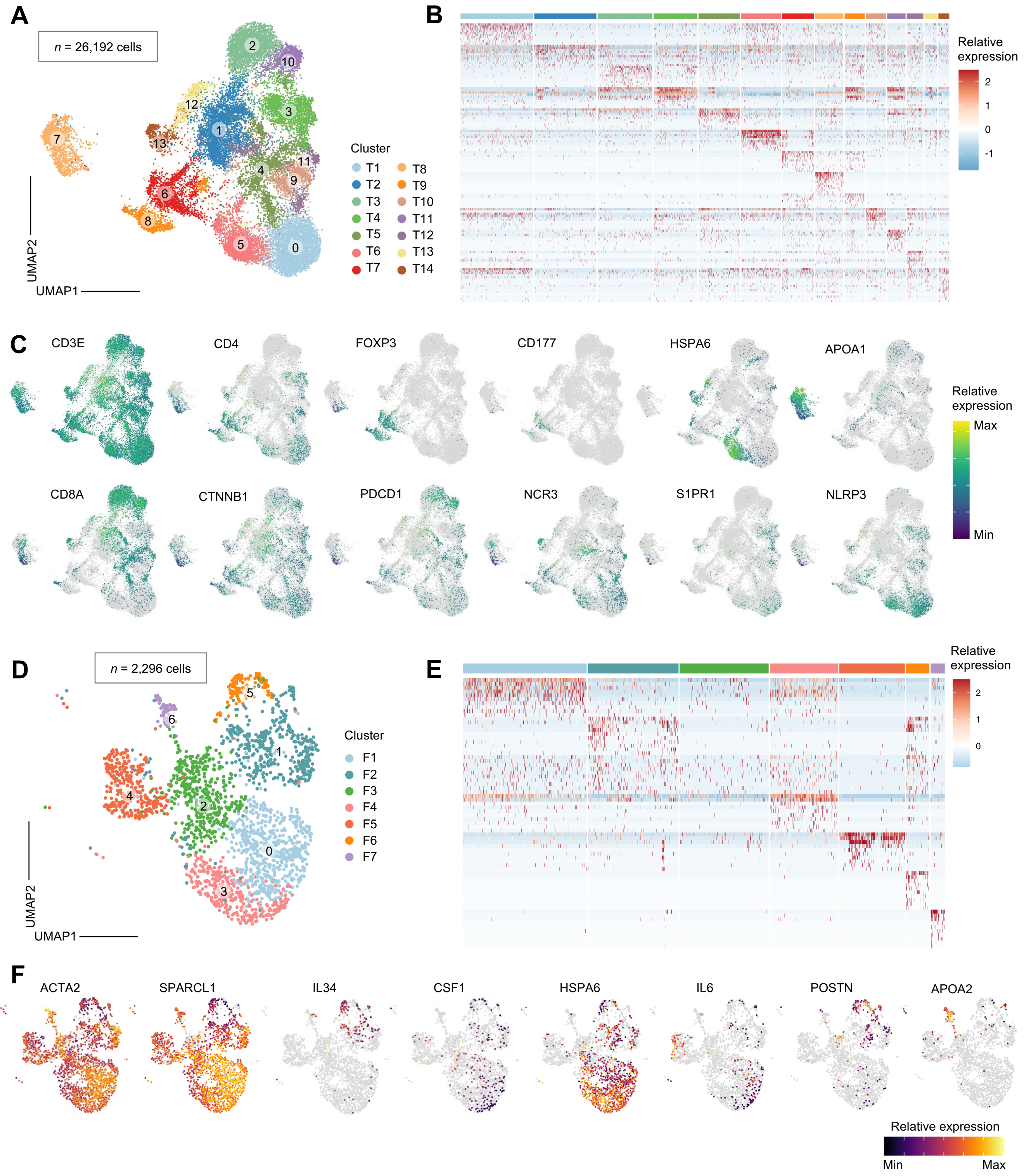


**Figure S3: scRNA-seq sub-clustering analyses of T cell and fibroblasts.**

**A.** UMAP showing 14 T cell clusters from 26,192 cells. **B.** Heatmap showing the top differentially expressed genes for the 14 T cell clusters. **C.** UMAP of selected genes representative of each cell type. **D.** UMAP showing 7 fibroblast clusters from 2,296 cells. **E.** Heatmap showing the top differentially expressed genes for the 7 fibroblast clusters. **F.** UMAP of selected genes representative of each cell type.

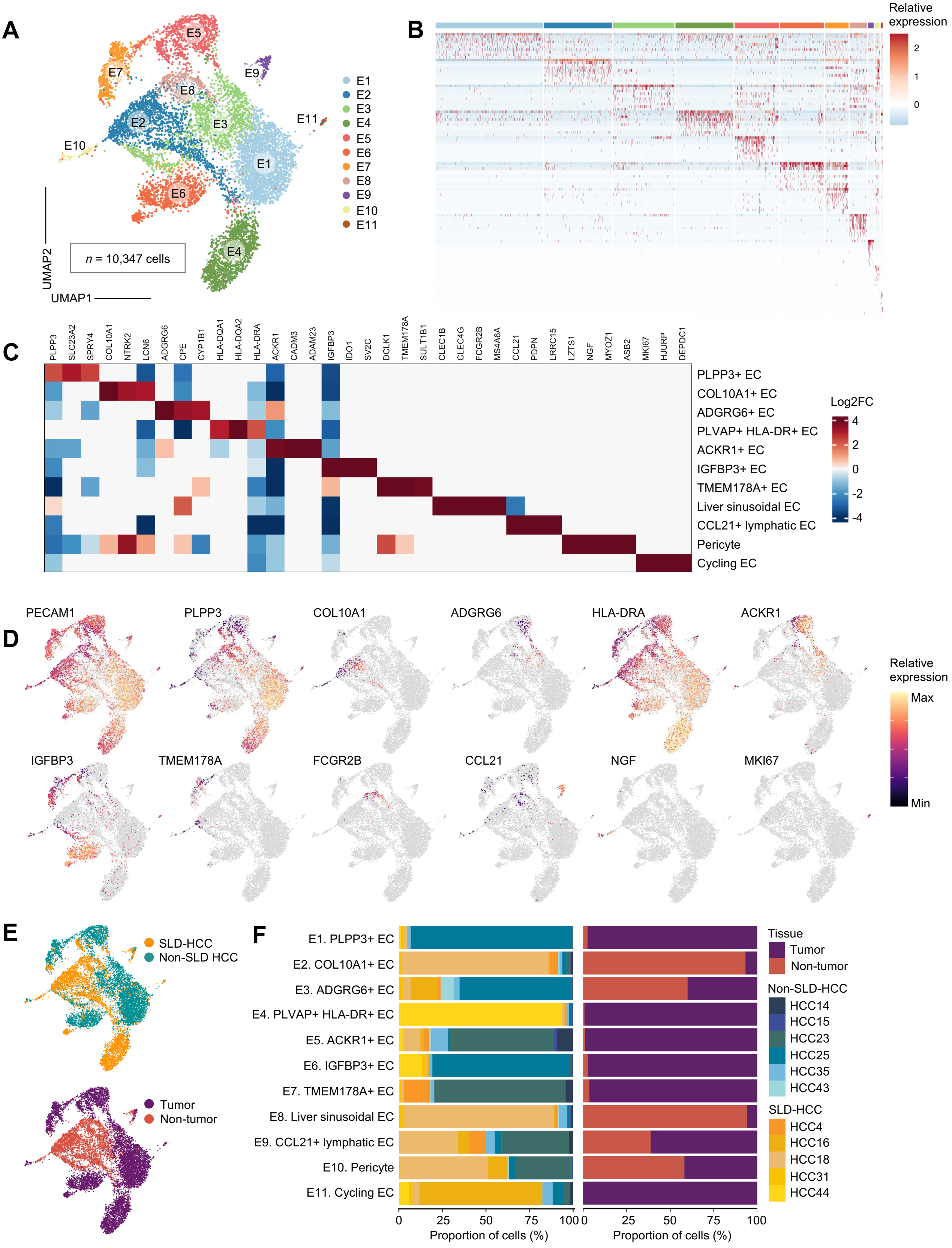


**Figure S4: scRNA-seq sub-clustering analysis of endothelial cells.**

**A.** UMAP showing 11 endothelial cell clusters from 10,347 cells. **B.** Heatmap showing the top differentially expressed genes for the 11 endothelial cell clusters. **C.** Heatmap showing relative expression of representative gene signatures for each sub-clusters. **D.** UMAP of selected genes representative of each cell type. **E.** UMAP showing distribution of cells according to disease status (upper) and tissue of origin (lower). **F.** Bar graph showing the proportion of each cell type in individual samples (left) and tissue of origin (right).

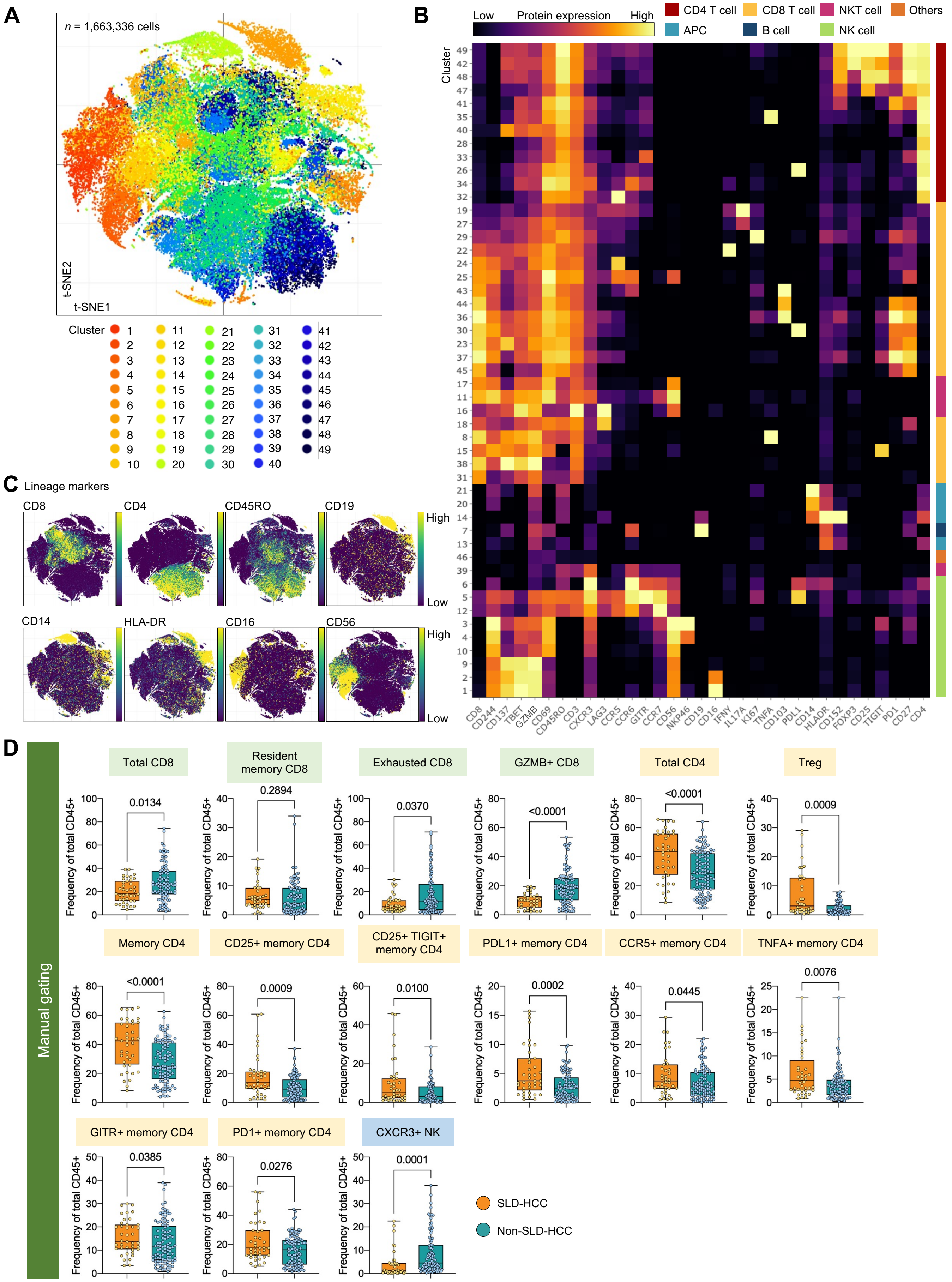


**Figure S5: Characterization of immune landscape in SLD- and non-SLD-HCC using CyTOF.**

**A.** t-SNE plot showing the distribution of 49 immune clusters according to their immune marker expression shown in **B**. **C.** t-SNE plots showing expression of selected immune markers **D.** The proportion of immune clusters in tumour tissues comparing SLD- and non-SLD-HCC by manual gating using FlowJo. Boxplots show median and the whiskers represent minimum and maximum values with the box edges showing the first and third quartiles. p-value determined by two-tailed Mann-Whitney test.

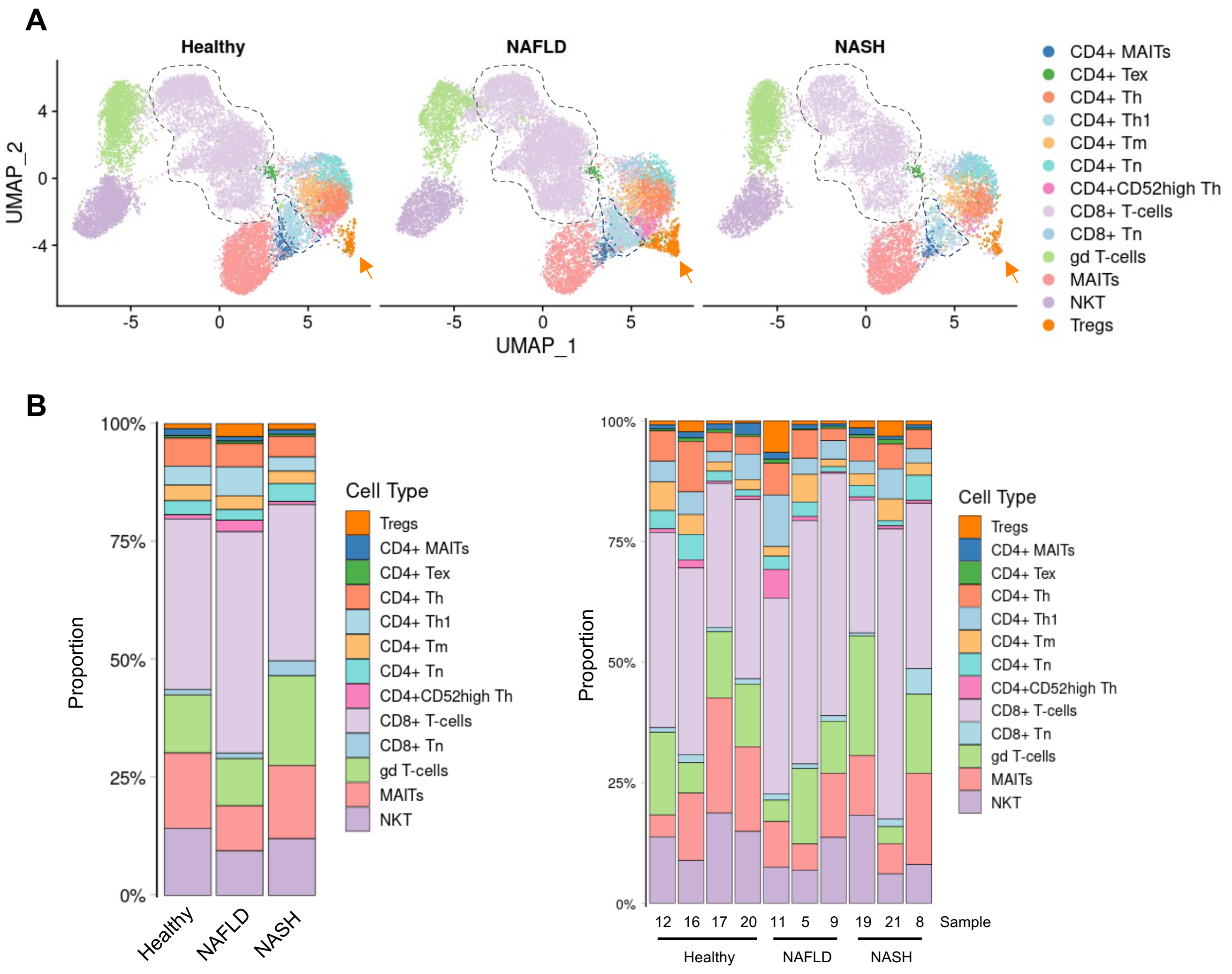


**Figure S6: scRNA-seq of T cells populations in healthy control, NALFD, and NASH patients.**

**A.** UMAP showing the distribution of T cell subsets in liver samples taken from healthy (n = 4), NAFLD (n = 3), or NASH (n = 3) patients. **B.** Bar graphs showing the proportion of T cell subsets in liver samples taken from healthy (n = 4), NAFLD (n = 3), or NASH (n = 3) patients.

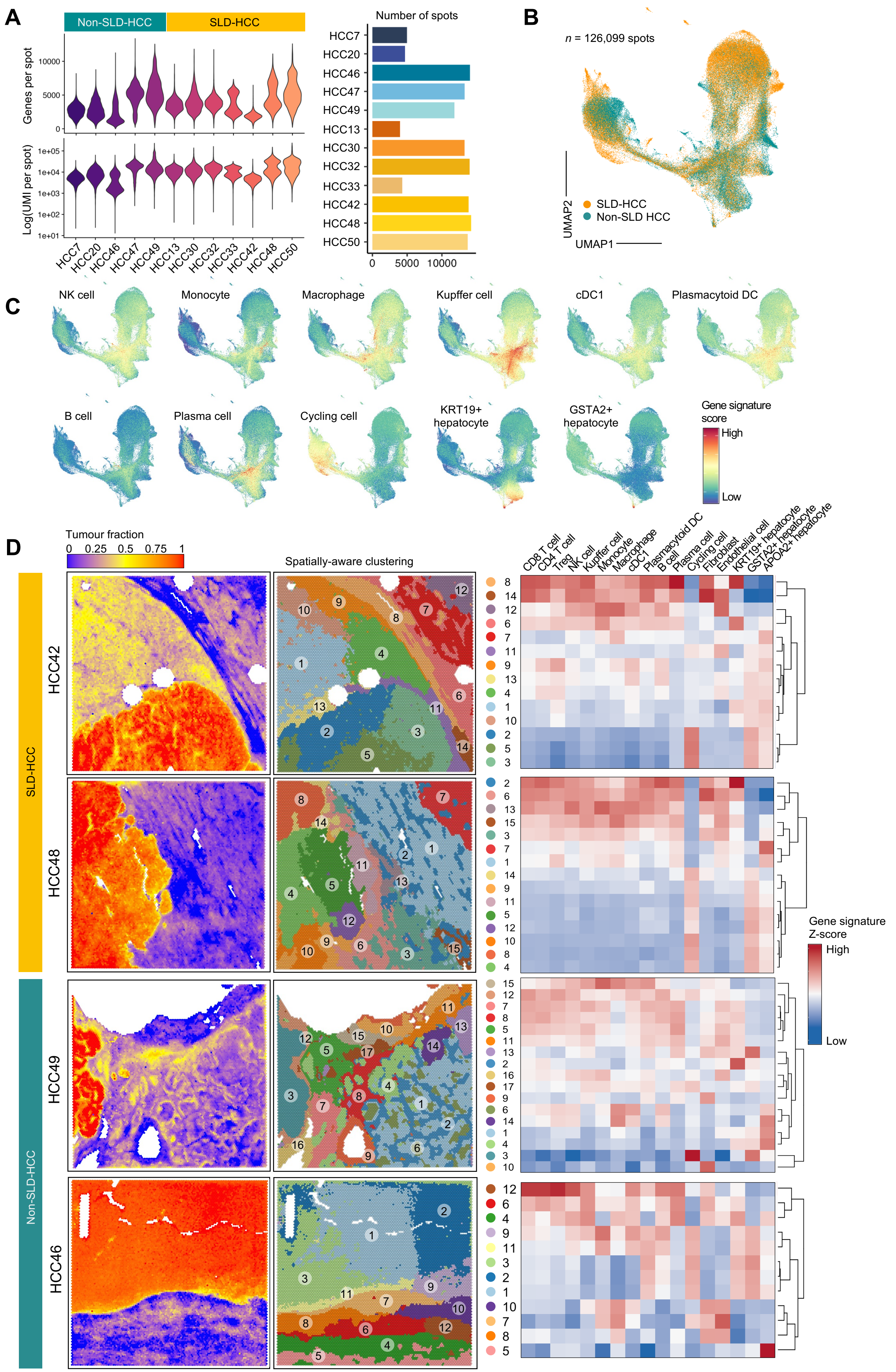


**Figure S6 (continued on the next page): Visium spatial transcriptomics data analysis on SLD- and non-SLD-HCC.**

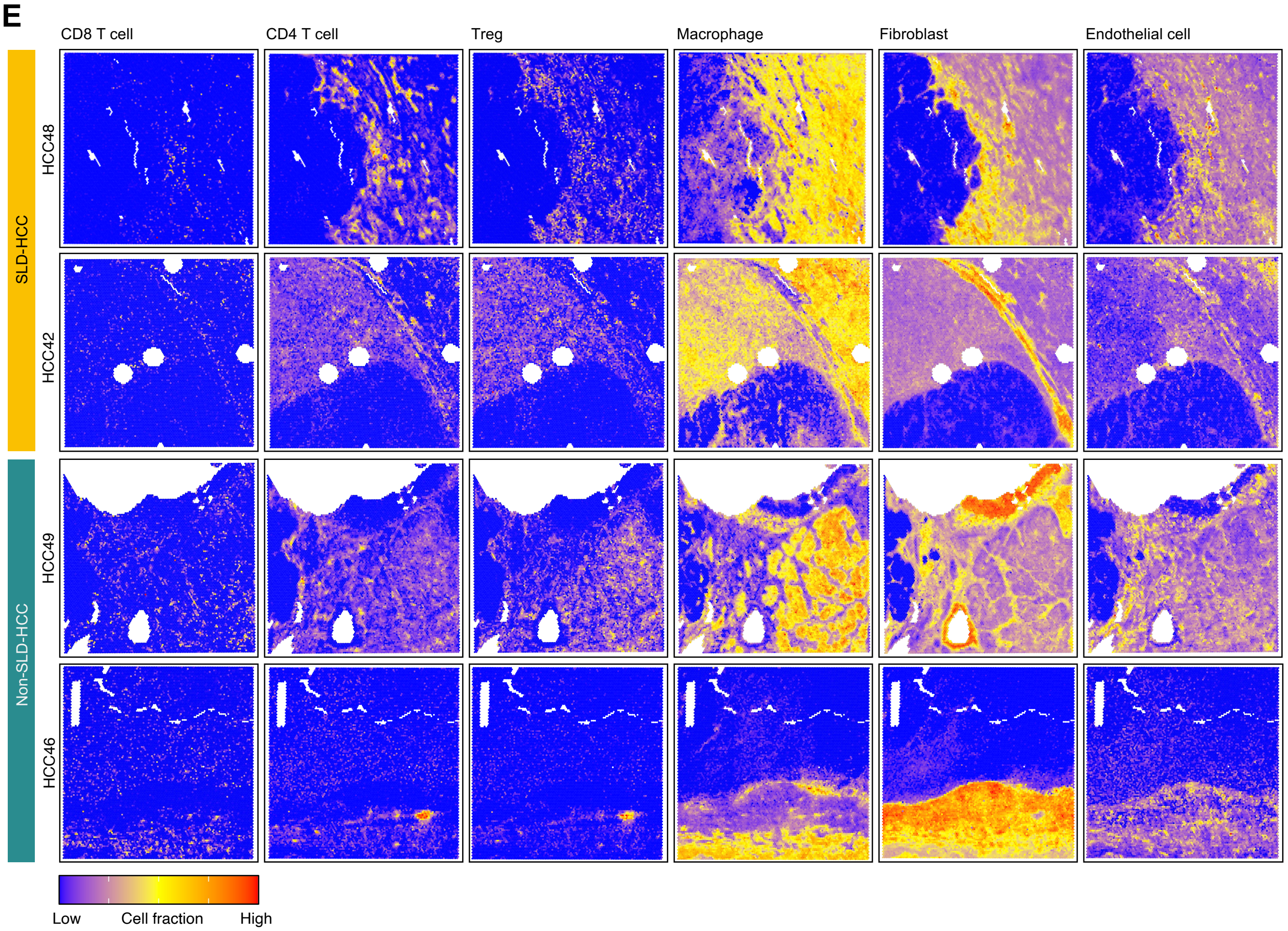


**Figure S7 (continued): Visium spatial transcriptomics data analysis on SLD- and non-SLD-HCCs.**

**A.** Violin plot showing the number of unique genes (upper) and unique transcripts (lower) per spatial spot, and the number of spots in each FFPE tissue sample analysed by Visium (10x Genomics) **B.** UMAP showing the distribution of Visium spatial data in SLD- and non-SLD-HCC. **C.** Scoring of cell types based on gene signatures from our scRNA-seq dataset on Visium. **D.** Representative images showing tumour identification using SpaCET (left) and spatially-aware clustering using Banksy showing the different domains within each tissue sample (middle), and the normalized gene signature scores for each cell type within the different domains (right) in SLD-HCC and non-SLD-HCC. **E.** Spatial deconvolution of selected cell types using SpaCET. **D and E.** Each field of view (FOV) = 11 x 11 mm.

**A**

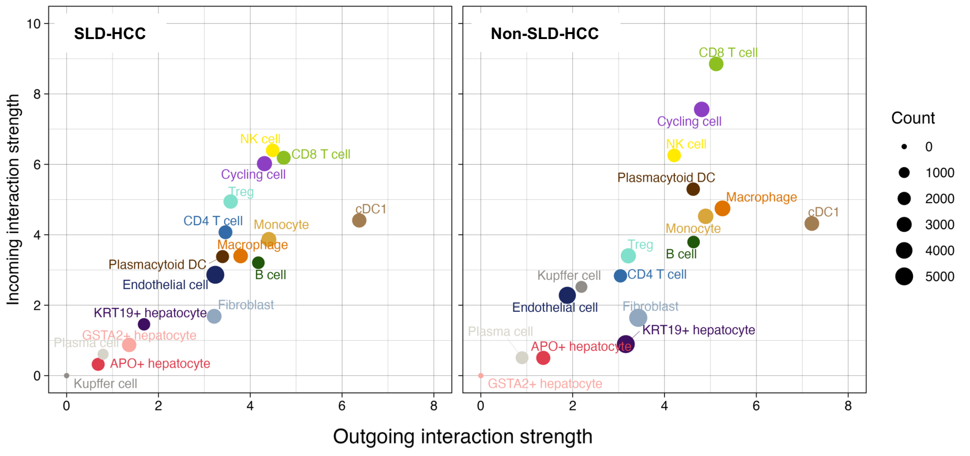


**Figure S8: Cell-cell interaction analysis using CellChat.**

**A.** CellChat analysis showing the incoming (receiver) and outgoing (sender) interaction signaling strength among all subsets based on scRNA-seq data.

A

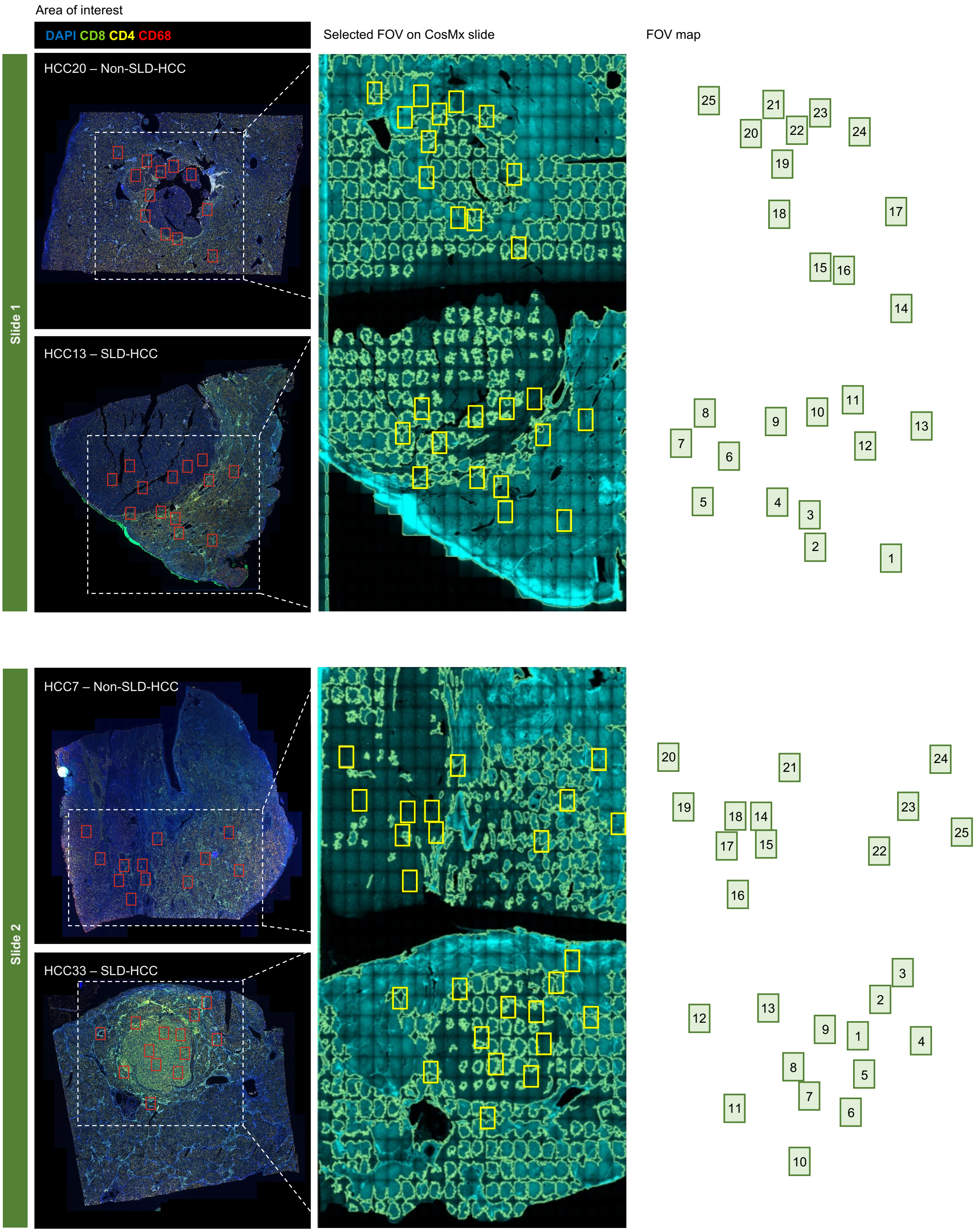


**Figure S9: CosMx spatial transcriptomics FOV selection.**

**A.** Overview for CosMx field of views (FOVs; n = 50) selection and data collection in two SLD- and two non-SLD-HCC. Adjacent tissue sections were stained for CD8, CD4, CD68, and nuclei (DAPI) and approximate area of interest indicated by red rectangles (left). FOVs selected on CosMx slides were indicated by yellow rectangles (middle) and the number of FOVs were mapped (right). Each FOV measures 0.7 x 0.9 mm.

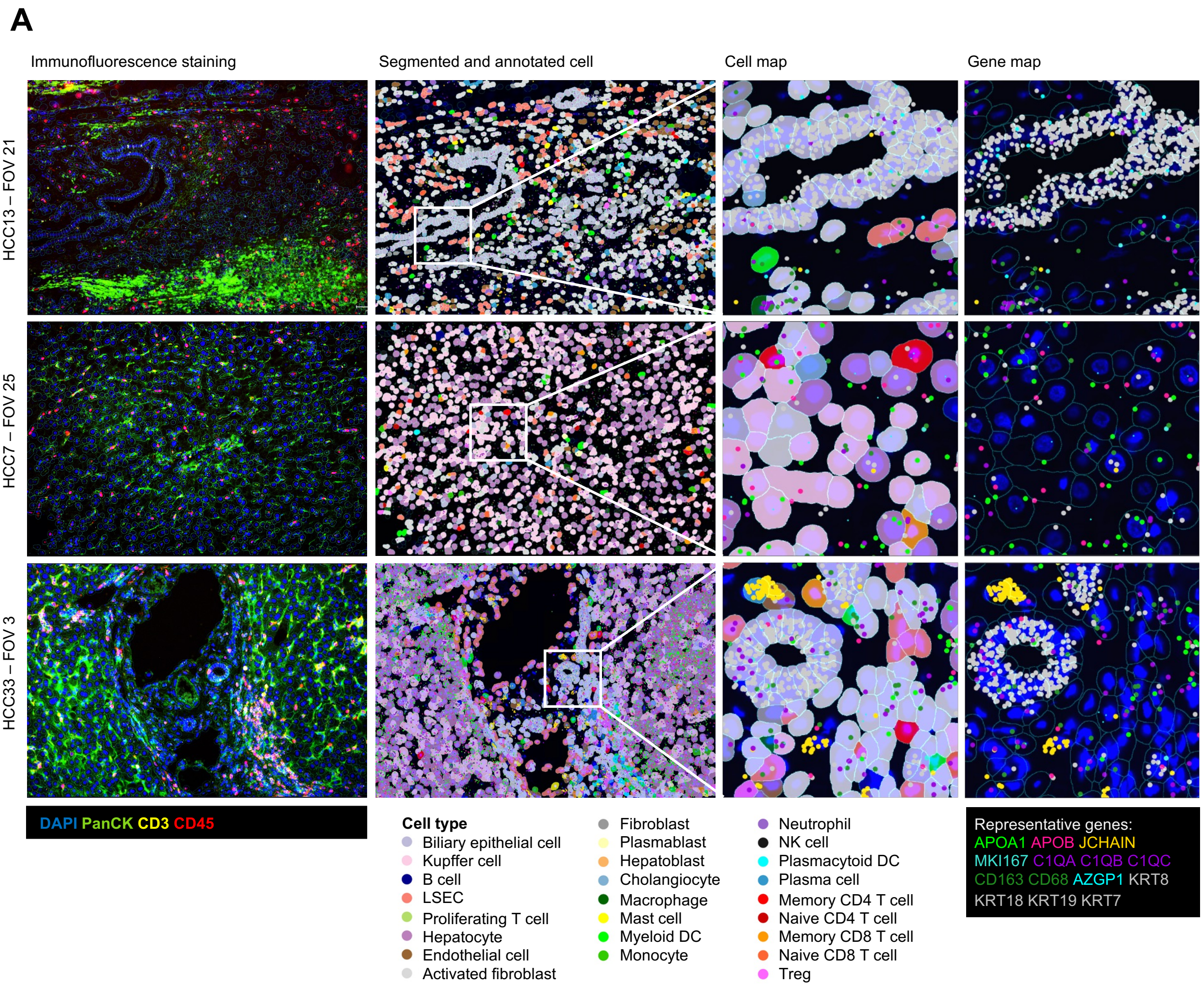


**Figure S10: Spatial transcriptomics analysis and clustering using CosMx.**

**A.** Representative CosMx filed of views (FOVs) showing immunofluorescence (IF) staining of PanCK (tumours marker), CD45 (Pan-immune marker), CD3 (pan-T cell marker) and DAPI (nuclear staining) along with cell segmentation and annotation based on selected marker genes (black box). Each subsets are indicated by colour code. Each FOV measures 0.7 x 0.9 mm.

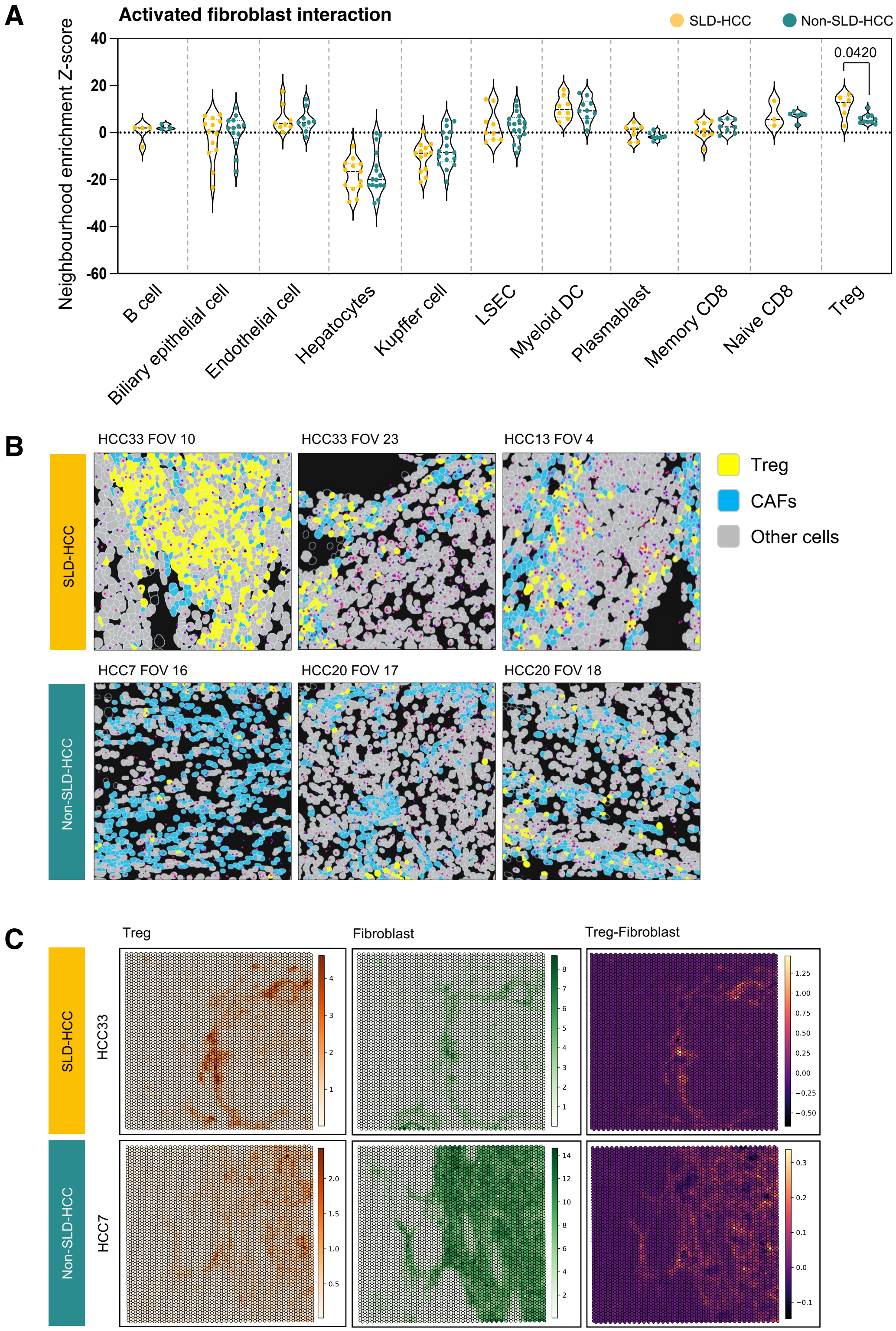


**Figure S11: Spatial transcriptomics cell-cell interaction analysis.**

**A.** Neighborhood enrichment score showing interaction strength between activated fibroblast and other cells in CosMx data. **B.** Representative CosMx field of views (FOVs) showing proximity between Treg and activated fibroblast in SLD- and non-SLD-HCC. Each FOV = 0.7 x 0.9 mm. **C.** Cellular deconvolution of Treg and fibroblast using Cell2location algorithm on the Visium spatial transcriptomics data based on gene signature from scRNA-seq. Images showing the distribution of Treg-fibroblast interaction in SLD- and non-SLD-HCC. Each FOV = 6.5 x 6.5 mm.

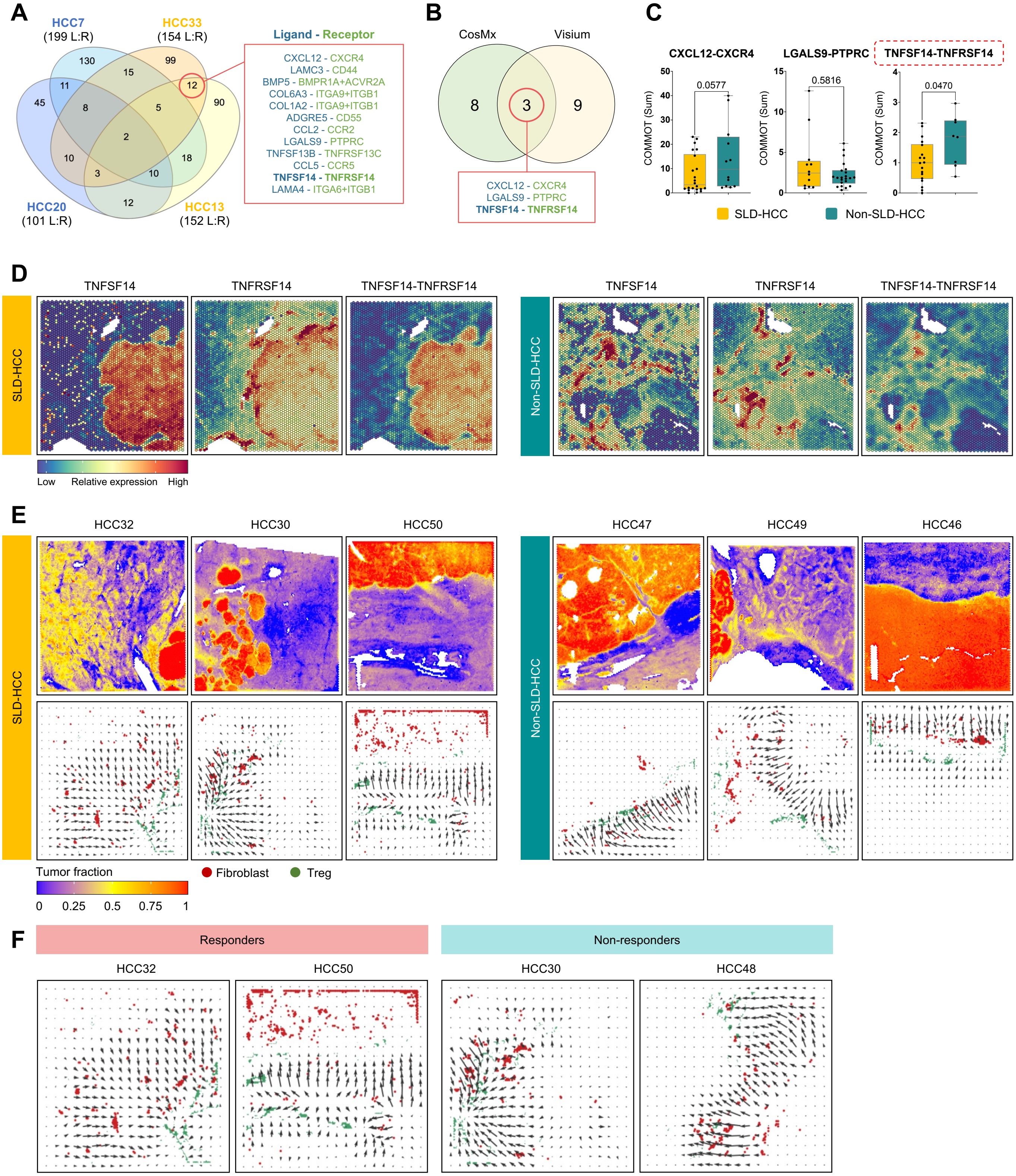


**Figure S12: Cell-cell interactome analysis by spatial transcriptomics.**

**A.** Comparison of LRs exclusively enriched in the margins of SLD-HCCs and non-SLD-HCCs. **B.** Mutual LRs identified using CosMx and Visium analysis. **C.** COMMOT scores comparing SLD- vs. non-SLD-HCCs of the 3 common LRs identified using CosMx and Visium. **D.** Expression of TNFSF14-TNFRSF14 on representative Visium data. Each field of view (FOV) = 6.5 x 6.5 mm. **E.** Tumor scoring and signalling of TNFSF14-TNFRSF14 between Treg and fibroblast from representative SLD- and non-SLD-HCCs analysed using COMMOT (Visium). **F.** Signalling of TNFSF14-TNFRSF14 between Treg and fibroblast from representative responders and non-responders to immunotherapy from SLD-HCC patients, analysed using COMMOT. **E, F.** The size and length of the arrows indicate strength and direction of the arrows indicates directionality of TNFSF14-TNFRSF14 interaction. Each FOV = 11 x 11 mm.
